# Flagellar switch inverted repeat impacts flagellar invertibility and varies *Clostridioides difficile* RT027/MLST1 virulence

**DOI:** 10.1101/2023.06.22.546185

**Authors:** Nguyen T. Q. Nhu, Huaiying Lin, Ying Pigli, Jonathan K. Sia, Pola Kuhn, Evan S. Snitkin, Vincent Young, Mini Kamboj, Eric G. Pamer, Phoebe A. Rice, Aimee Shen, Qiwen Dong

**Author notes:** Address correspondence to Qiwen Dong,. Nguyen T. Q. Nhu, Division of Infectious Diseases, Department of Internal Medicine, University of Michigan, Ann Arbor, Michigan, USA; Qiwen Dong, Department of Molecular Biology and Microbiology, Tufts University, Boston, Massachusetts, USA.

## Abstract

*Clostridioides difficile* RT027 strains cause infections that vary in severity from asymptomatic to lethal, but the molecular basis for this variability is poorly understood. Through comparative analyses of RT027 clinical isolates, we determined that isolates that exhibit greater variability in their flagellar gene expression exhibit greater virulence *in vivo*. *C. difficile* flagellar genes are phase-variably expressed due to the site-specific inversion of the *flgB* 5’UTR region, which reversibly generates ON vs. OFF orientations for the flagellar switch. We found that longer inverted repeat (IR) sequences in this switch region correlate with greater disease severity, with RT027 strains carrying 6A/6T IR sequences exhibiting greater phenotypic heterogeneity in flagellar gene expression (60%-75% ON) and causing more severe disease than those with shorter IRs (> 99% ON or OFF). Taken together, our results reveal that phenotypic heterogeneity in flagellar gene expression may contribute to the variable disease severity observed in *C. difficile* patients.

## INTRODUCTION

*Clostridioides difficile* (*C. difficile*) is the leading cause of hospital-acquired infection in the U.S., with an estimated incidence of approximately 224,000 cases per year.^1^ The incidence of community-acquired *C. difficile* infection (CDI) is also increasing, with up to 51.2 cases /100,000 population.^2^ Major risk factors for developing CDI include broad-spectrum antibiotic usage, as antibiotics deplete the commensal bacterial species that provide colonization resistance against CDI. However, CDI disease severity in humans ranges from asymptomatic colonization to diarrhea to severe pseudomembranous colitis and even death. Recent studies have revealed that the gut microbiota and host immunity impact CDI pathogenicity,^3^ but genetic features of *C. difficile* likely also regulate disease severity in humans.

*C. difficile* virulence requires the production of at least one of its glucosylating toxins, TcdA and/or TcdB.^4^ These toxins are internalized by epithelial cells via endocytosis^5–7^ and released into the cytosol, where they glucosylate and inactivate intracellular GTPases.^8–10^ These activities disrupt cell signaling pathways and cytoskeletal structure, leading to cell death.^11^ The impact of TcdA and TcdB on disease progression and severity, however, can vary, depending on the toxin titer, the strain of *C. difficile*, and the host’s innate and adaptive immune defenses.^12–14^

In addition to toxin expression, flagellar-mediated motility has been suggested to modulate *C. difficile* virulence. Flagellar motility contributes to the virulence of *S. enterica* and *E. coli* by enhancing host cell invasion, adhesion to epithelial cells, and systemic inflammation.^15–17^ However, the impact of flagellar motility on *C. difficile* infection *in vivo* remains controversial and is likely strain- and host-dependent.^18–20^ Deletion of *fliC*, which encodes flagellin, did not impact the virulence of *C. difficile* R20291 in a germ-free mouse model, yet it reduced its virulence in antibiotic-treated SPF mice.^18,19^ In contrast, the deletion of *fliC* in *C. difficile* strain CD630 increased its virulence in a hamster model of CDI.^20^ Notably, flagellar genes are heterogeneously expressed in *C. difficile* due to the inversion of a flagellar switch sequence by the tyrosine recombinase RecV.^21^ Inversion of this region in the 5’UTR of *flgB* to the ON orientation allows for expression of a flagellar gene operon that includes the gene encoding the sigma factor, SigD, which directly induces the expression of flagellar gene operons as well as *tcdR*, which encodes another sigma factor that activates toxin gene expression.^22^ Thus, flagellar gene expression is coupled with toxin gene expression.^21,23^ Despite these insights, the impact of *C. difficile* flagella and the heterogeneity of the flagellar gene expression on CDI virulence remains unclear.

In this study, we investigated the relationship between flagellar gene expression and virulence in *C. difficile* ST1 strains isolated from patients with CDI. Strains of the multi-locus sequence type 1 (ST1), also known as the NAP1/B1/ribotype 027, are highly transmissible, have increased multidrug resistance,^24,25^ and were initially reported to be hyper-virulent.^26,27^ However, heterogeneity in the virulence^28,29^ and toxin production of ST1 strains^30,31^ have been reported, revealing that there is considerable variation between individual ST1 trains. By examining 22 ST1 clinical isolates in a mouse model of infection, we observed marked differences in virulence. We found that sequence differences in the flagellar switch region, specifically in the inverted repeat (IR) region, of these ST1 strains correlated with disease severity. Our data reveal that these sequence differences alter the invertibility of the flagellar switch, with longer IR sequences causing greater heterogeneity within the population, i.e., a mixture of flagellar gene ON and OFF, and shortened IR sequences being associated with more fixed populations of either flagellar gene ON or OFF. Furthermore, exchanging a longer IR sequence with a shorter IR sequence in *C. difficile* R20291 significantly reduced its virulence, particularly when combined with the OFF flagellar switch orientation. Our results argue that flagellar switch IR sequences alter the invertibility of the flagellar switch, which contributes to the considerable heterogeneity in virulence observed between *C. difficile* ST1 strains.

## RESULTS

### The flagellar expression associates with the virulence of clinical *C. difficile* RT027 isolates

We previously showed that the *in vivo* virulence of *C. difficile* ST1 isolates in C57BL/6 mice varies widely, ranging from mortality to the absence of any weight loss or diarrhea.^31^ While we showed that the avirulence of 2 ST1 strains was due to a small internal deletion in the *cdtR* gene, the remaining ST1 strains did not exhibit any genetic variability in their pathogenicity and CDT loci, which encode the TcdA and TcdB glucosylating toxins and the binary toxin CDT, respectively^31^ (**Figure S1A**). In accordance, the virulence differences (% weight loss) observed between these strains did not correlate with the fecal toxin levels measured using a cell-based toxicity assay (**Figures S1B-C**). In addition, no correlation in the colonization (CFU) levels on day 1 post-infection and virulence of the strains was observed (**Figure S1B-S1D**). To identify additional mechanisms for variable virulence among ST1 strains, we selected two isolates that induced >10% weight loss (ST1-12 and ST1-53) and two isolates that resulted in < 10% weight loss (ST1-6 and ST1-27) for further study (**Fig. S1**). Here, we refer them as high-virulence and low-virulence isolates.

We first compared the transcriptional profile of the four isolates during infection of antibiotic-treated mice by harvesting cecal contents one-day post-infection and conducting RNAseq profiling of the isolates (**Figure 1A**). The transcriptomic profile revealed that flagellar-related genes are over-expressed in high-virulence isolates relative to low-virulence isolates (**Figure 1B**). These differences in flagellar gene expression were also observed when RT-qPCR was used to compare the expression of two flagellar genes, *fliE* and *fliS1*, between the strains using the same cecal content samples (**Figure S2A**). High-virulence isolates also exhibited greater flagellation and spread further on swim plates 24-hours after inoculation (**Fig. 1C-1D)**. In contrast, the low-virulence isolates were aflagellate and exhibited delayed spreading on the swim plates (**Fig. 1C-1D)**. Notably, the RNA-Seq analyses confirmed that toxin genes, including *tcd* genes and *cdt* genes, were expressed at similar levels between the four strains during murine infection (**Fig. 1B**), consistent with the similar levels of fecal toxins detected during infection in mice. While ST1-27 appeared to colonize mice at higher levels compared to the other three isolates (**Fig. S2B-S2C**), the growth of these 4 isolates did not differ from each other in broth culture (**Fig. S2D**). Interestingly, according to the RT-qPCR results performed with the same cecal content samples, the high-virulence strain ST1-12 expressed significantly higher *tcdA* and *tcdB* relative to the other three strains, including the other high-virulence strain ST1-53. Though it did not fully reproduce the RNAseq observation, it suggests again that toxin production differences cannot fully explain the variation in virulence observed among ST1 isolates (**Figure S2A**).

**Figure 1:**
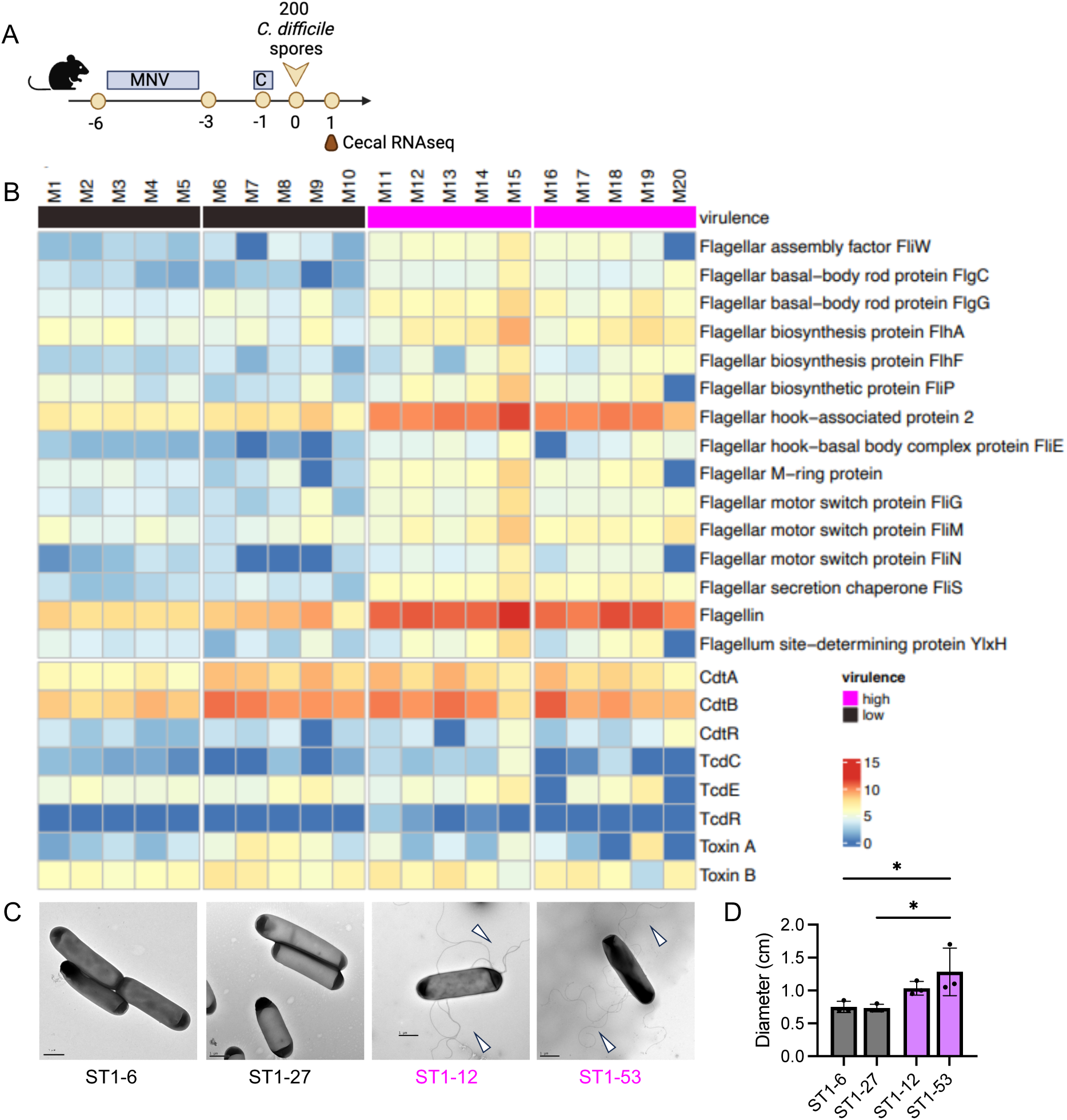
High virulence isolates make more flagella. (A) The schematic of the mouse infection. (B) Heat map of RNA sequencing results from the cecal contents of mice infected with the four ST1 isolates shown. The heat map is generated based on the number of transcripts per sample after normalization. RNA was extracted from the cecal contents of mice one-day post-infection (n= 5 mice per isolate). (C) Transmission electron micrographs of the four isolates. (D) Swim plates results of four isolates after 24 hours incubation (n = 3 replicates per isolate). High-virulence isolates: pink. Low-virulence isolates: black. Statistical significance was calculated by One-way ANOVA, * p < 0.05.

### Variation in the flagellar switch inverted repeats of RT027/ST1 clinical isolates

The difference in flagellar gene expression between high-virulence and low-virulence isolates led us to compare their flagellar genomic regions. While the flagellar genes were identical between the isolates, we identified variation in the flagellar switch region, which plays an important role in regulating the expression of the flagellar operons.^21^ The flagellar switch region comprises a central region flanked by a left-inverted repeat (LIR) and a right-inverted repeat (RIR) (**Figure 2A**). The inverted repeats are recognized by the tyrosine recombinase RecV, which inverts the central region. This leads to the switch region exhibiting either an ON or OFF orientation, which allows or inhibits flagellar gene expression, respectively ^21^. When we expanded the analysis of the flagellar switch region to all 68 ST1 isolates in our collection, we found that 65 of the isolates have a flagellar switch central sequence identical to the reference RT027/ST1 strain, R20291, and thus carried for further analyses. Most notably, we found variability in the sequence of the left and right IRs that flank the central flagellar switch region. Specifically, while most isolates have IRs identical to the reference strain *C. difficile* R20291, 40% of isolates have at least one less A or T in either the left or right IRs (**Figure 2A and Table S1**). In total, four types of IR flanking regions were identified in our strain collection; we named them: Common1 (C1-59.38% 6A/6T), Common2 (C2-32.81%, 6A/5T-ON or 5A/6T-OFF), Rare1 (R1-6.25%, 5A/6T-ON or 6A/5T-OFF), and Rare2 (R2-1.56%, 5A/5T). The C1 switch region, the most common sequence, is identical to that observed in *C. difficile* R20291, with 6 A’s on the left IR (LIR) and 6 T’s on the right IR (RIR). When C1 inverts between the OFF vs. ON orientation, the number of A’s and T’s in both IRs remains the same. In contrast, the composition of the Common 2 (C2) region differs depending on the orientation of the switch region: in the OFF orientation, the LIR consists of 5 A’s and the RIR consists of 6 T’s (C2-OFF); in the ON orientation, the LIR consists of 6 A’s and the RIR consists of 5 T’s (C2-ON). The Rare 1 region (R1) also has an asymmetric distribution depending on the switch orientation except that in the OFF orientation, the LIR consists of 6 A’s and the RIR consists of 5 T’s (R1-OFF). The Rare 2 region (R2) IRs remain the same regardless of the switch orientation, with the LIR consisting of 5 A’s and the RIR consisting of 5 T’s. However, the R2 is only observed in one strain, ST1-67, where it is in the OFF orientation. Notably, Sanger sequencing of the flagellar switch region confirmed the HiSeq whole genome sequencing analyses (**Table S1**).

**Figure 2:**
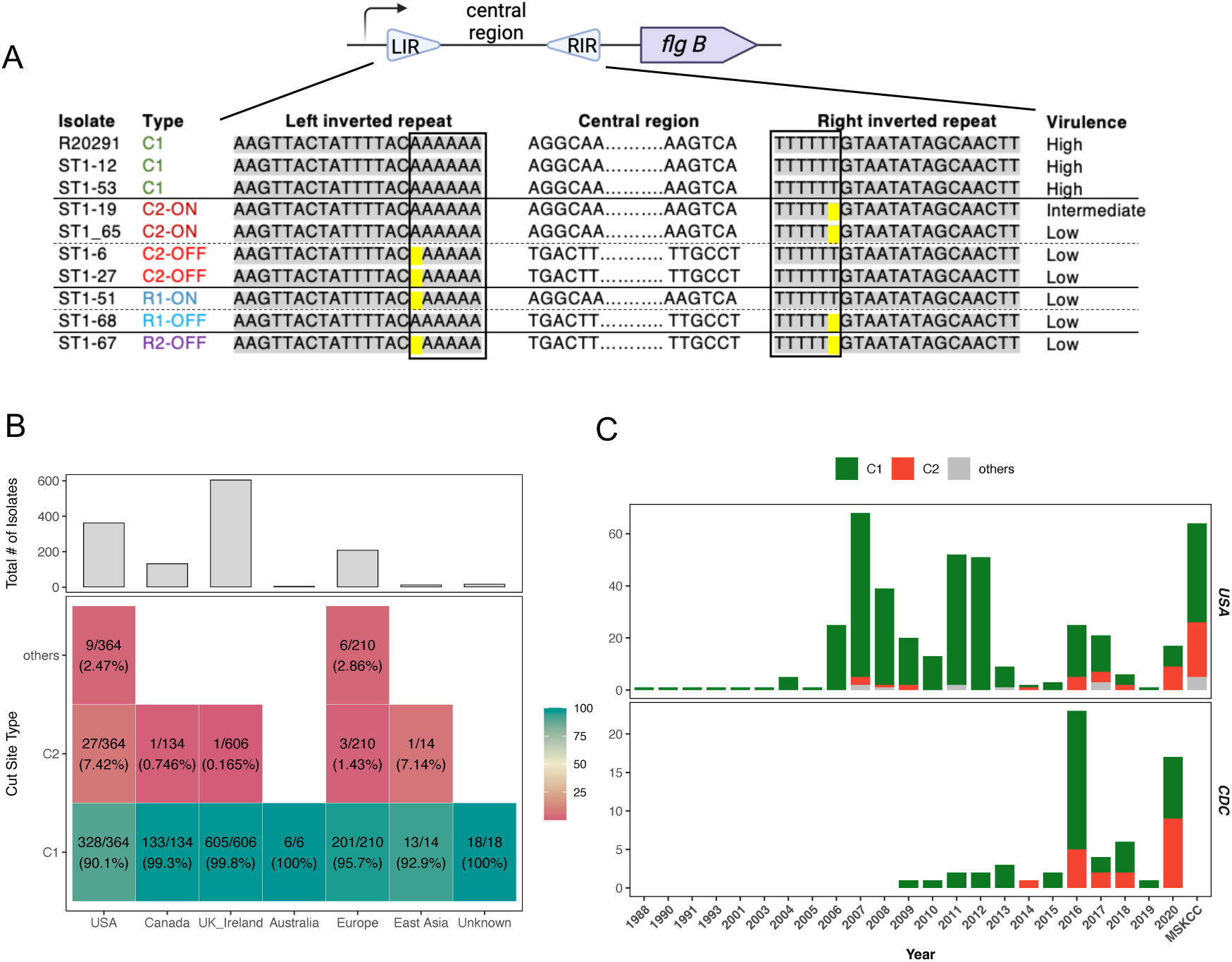
Variations in flagellar inverted repeat types observed among ST1 isolates. (A) Schematic of the flagellar switch and the alignment of the flagellar switch regions from representative isolates. In Illumina sequencing the flagellar switch region of the Common 1 IR type (C1, 6A/6T) is observed in the ON orientation. The Common 2 IR type (C2, 6A/5T) is observed in isolates with the flagellar switch region being both in the ON and OFF orientations. Rare 1 (R1-5A/6T) was observed in isolates with the flagellar switch being ON and OFF. The Rare 2 IR type (R2, 5A/5T) was observed only in one isolate with the flagellar switch region being in the OFF orientation. Full list is in Table S1. (B) IR types of isolates collected from around the world. (C) IR types of isolates collected in the US and by the CDC.

To assess the conservation of the IR types observed, we screened 1,359 RT027/ST1 isolates whose whole-genome sequences were available from NCBI BioSamples. These isolates derive from the CDC HAI-Seq *C. difficile* collection (PRJNA629351)^32^ and Texas and Michigan medical centers (PRJNA595724, PRJNA561087).^33^ While isolates from more recent publications were also included,^34–37^ all *C. difficile* isolates were collected from 1988 to 2020. Most isolates (1318/1359) derive from Continental Europe, UK, Ireland, and the U.S., with a small number (20/1359) of isolates from East Asia and Australia. Most *C. difficile* ST1 isolates carry the flagellar switch IR type C1, representing 96.5% (1311/1359) of isolates analyzed. Type C2 was observed in only 2.4% (33/1359) of isolates, of which most (27/33) were obtained in the U.S between 2007 and 2020. The other IR types were detected at even lower frequencies, representing only 0.66% of isolates (**Figure 2B**).

When more recent CDC HAI-Seq collection strains were analyzed, which were predominantly submitted between 2009 and 2020, the IR type C1 remained dominant, but the C2 represented 30% (19/63) isolates, similar to our collection (32.8%). Interestingly, the proportion of C2 increased in isolates collected in 2020, accounting for 53% (9/17) of isolates collected in 2020 (**Figure 2C**). Thus, our data suggest that variability in *C. difficile* flagellar IR regions has increased over time, although their impact on *C. difficile* virulence remains unclear.

### Flagellar invertibility has a strong association with virulence of RT027/ST1

To analyze the impact of inverted repeat types on RT027/ST1 virulence, we correlated the weight loss in mice caused by *C. difficile* infection with different IR types. We found a strong correlation between IR type C1 and more severe weight loss in infected mice (p = 3.2e-05) (**Figure 2A and 3A**). Since a small deletion in the RIR and surrounding region greatly reduces the invertibility of *C. difficile* flagellar switch, essentially locking it in one orientation,^23^ we tested whether the different IR types impact the invertibility of the flagellar switch region. Using orientation-specific qPCR analyses to quantify the fraction of ON and OFF cells in *C. difficile* population from liquid culture (**Figure 3B**),^21^ we found a weak correlation between greater weight loss and strains with a higher proportion of ON cells in a population by linear regression (R^2^ = 0.39). However, the most virulent isolates had ∼70 % of cells in the ON orientation and consisted of the C1 IR type (**Figure 3C**, green circles). Furthermore, isolates with the other IR types exhibited lower virulence than the C1 isolates and were either nearly 100% ON or 100% OFF (**Figure 3C**). Thus, our results suggest that IR type governs the flexibility of the flagellar switch region and that IR types that lead to more heterogeneous flagellar gene expression, i.e. the C1 IR type, enhance the virulence of *C. difficile* strains.

**Figure 3:**
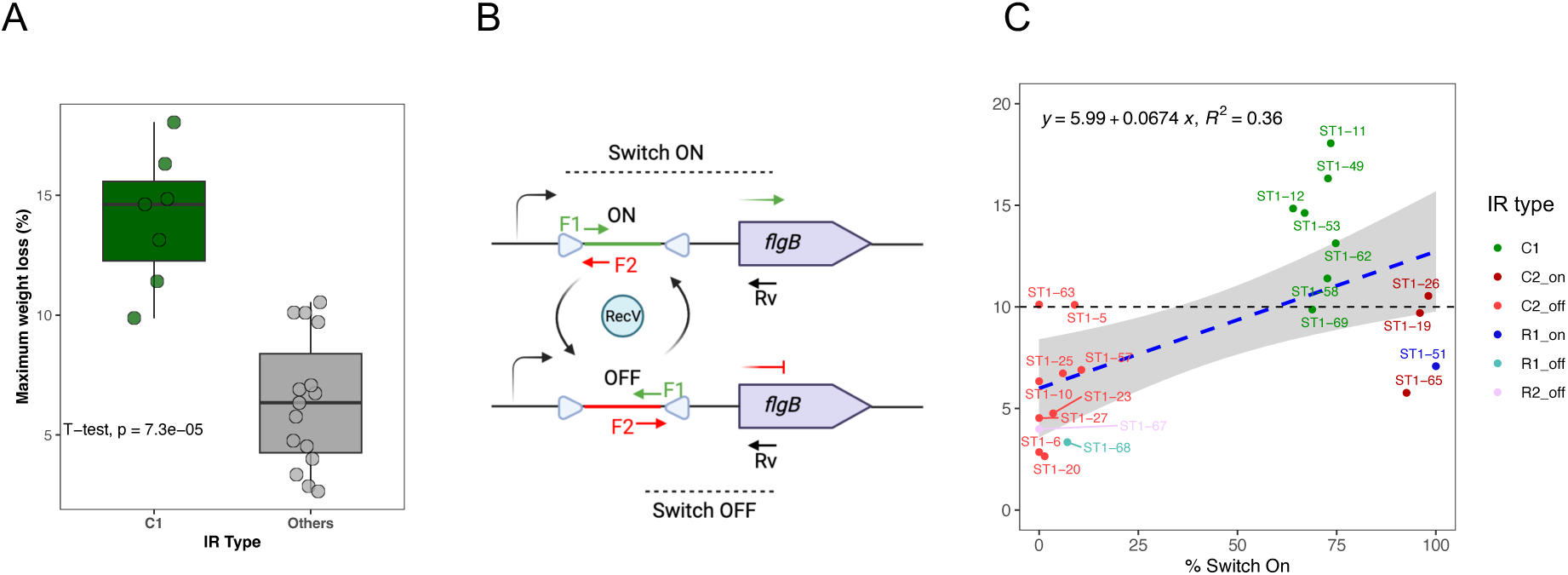
High virulence isolates exhibit more heterogeneous flagellar gene expression. (A) %maximum weight loss graphed by IR types. (B) Schematic of the qPCR strategy to measure the proportion of flagella - ON vs. OFF cells in the population. (C) Correlation between % maximum weight loss and % flagella – ON cells. DNA was extracted from bacterial cultures during logarithmic growth.

### The Inverted repeat type correlates with the invertibility of the flagellar switch

To further investigate whether different IR types impact the flagellar switch invertibility, analyzed the flagellar orientation of our 64 ST1 isolates across 3-4 biological replicates per isolate. We cultured individual isolates in broth to exponential growth and prepared the genomic DNA for orientation-specific qPCR analyses. When calculating the proportion of C1 cultures in the ON orientation, we found that the C1 IR type is slightly biased toward the ON orientation, with approximately 73% (55-99%) in the ON orientation (**Figure 4A**). In contrast, the biased-OFF isolates, including C2-OFF, R1-OFF and R2-OFF, have 95.6% (88.7-100.0) of their population in the OFF orientation, whereas the biased-ON isolates, including C2-ON and R2-ON, have 96.7% (92.6-100.0) of their population in the ON orientation (**Figure 4A**). Such a strong correlation between IR types and biased-flagellar orientations indicates that mutations in the IR regions greatly impact the switching invertibility.

**Figure 4:**
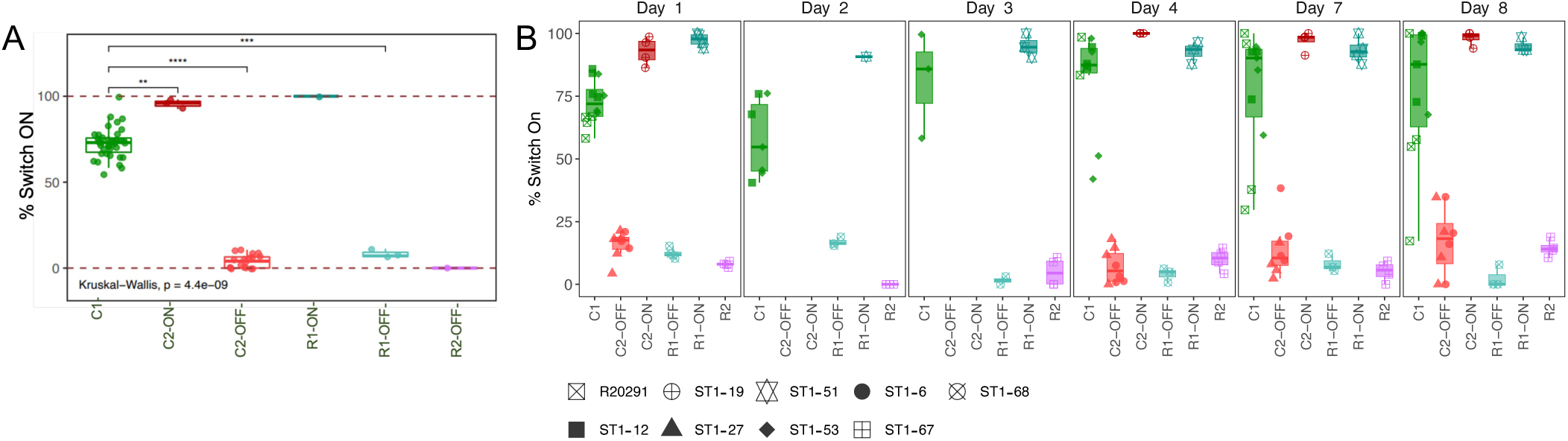
Flagellar inverted repeat types are associated with flagellar switch invertibility in culture and in mice. (A) Proportion of the population with the flagellar switch region in the ON orientation during logarithmic growth in broth culture and (B) during infection over time as determined by qPCR. The results reflect samples obtained from the feces (day1-7) and cecum (day 8) of *C. difficile-* infected mice. Statistical significance was calculated by One-way ANOVA, ** p< 0.01,, *** p < 0.001, **** p < 0.0001.

To test if the biased IR types also restrict the invertibility of *C. difficile* isolates *in vivo*, we infected mice with *C. difficile* R20291 and clinical isolates with different IR types and monitored the ON and OFF cell fractions in fecal or cecal contents over the course of eight days. On day 1 post-infection, we found that *C. difficile* isolates with the C1 IR type had about ∼75% of their population with the ON orientation (**Figure 4B**), similar to our broth culture analyses (**Figure 4A**). Interestingly, the proportion of C1 IR type strains in the ON vs. OFF cells exhibited greater variability over the course of the infection compared to the isolates with IR types C2, R1, and R2. These latter three IR type strains maintained their flagellar switch region in a biased orientation over the eight-day infection course (**Figure 4B and S3**). These data further highlight the strong correlation between IR type and flagellar switch invertibility. Specifically, the C1 IR type (6A/6T) exhibits greater invertibility in its flagellar switch region, while the other IR types appear to “lock” the flagellar switch in either the ON or OFF orientation.

### *C. difficile* flagellar IR type impacts RecV-mediated DNA inversion

Since IR types correlate with the invertibility of ST1 isolates (**Figure 4**), we wondered if the IR types specifically affect RecV-mediated DNA inversion. To test this possibility, we assessed the invertibility of IR types in a heterologous host using an *E. coli*-based colorimetric assay.^38^ *E. coli* were co-transformed with plasmids encoding *C. difficile recV* and a second plasmid containing a promoter flanked by either the C1, C2 or R2 IR type. Depending on the orientation of the promoter within the switch region flanked by the IR variants, it will drive the expression of either *gfp* or *rfp*. Thus, two variations of the reporter plasmid were transformed into *E. coli* for each IR type variant: One with the promoter initially driving *gfp* (left panel), and the second with the promoter driving *rfp* (right panel). RecV-mediated DNA inversion of the reporter plasmid results in a switch from red to green (or green to red) (**Figure 5A)**. When the reporter plasmid carried the C1 IR type, a high percentage of dual plasmid-transformed colonies exhibited a color switch (**Figure 5B).** In contrast, when the reporter plasmid carried the C2 IR type, only 1-2 colonies exhibited a color switch, whereas no colonies were observed to have color-switched when the reporter plasmid carried the R2 IR type (**Figure 5B**). These results align with our qPCR analyses of the switch region, strongly suggesting that the IR sequence controls the invertibility of the flagellar switch region.

**Figure 5:**
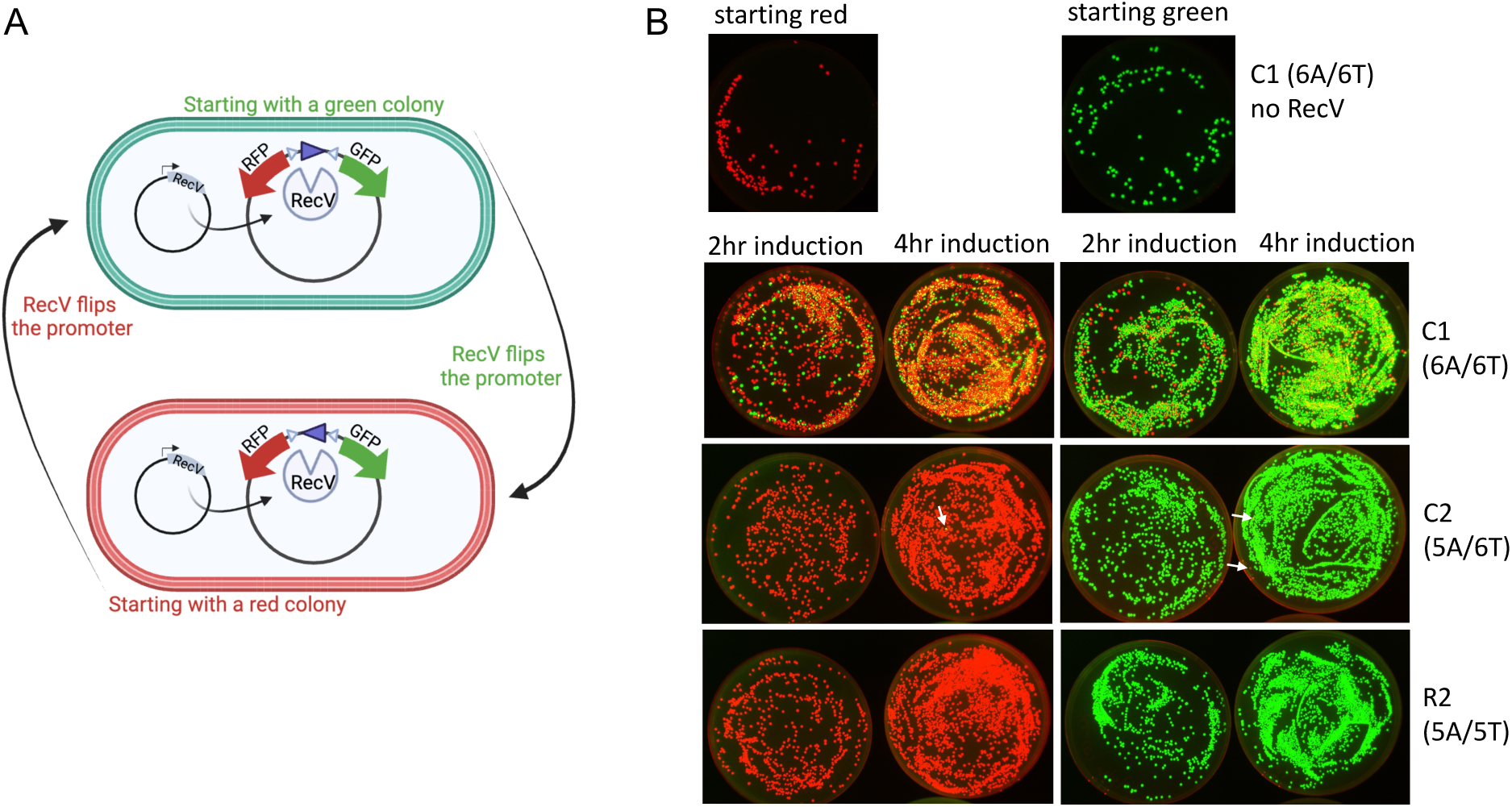
Flagellar inverted repeat (IR) types determine the flagellar switch invertibility. (A) Schematic of the color switch assay in *E. coli*. Two plasmids were transformed into *E. coli.* One plasmid encodes *C. difficile* RecV. The other plasmid contains an invertible sequence where the promoter is flanked by the inverted repeat (IR) variants. Depending on the orientation of the promoter, either *gfp or rfp* will be expressed, and color switch from red to green (or green to red) is expected upon RecV acting on flexible switches. (B) Color switching assay when different types of flagellar inverted repeats flank the invertible region. The switch frequency of the Common 1 (C1-6A/6T), Common 2 (C2-5A/6T), Rare 2 (R2-5A/5T) IR variants was visualized. White arrows point to colonies that underwent a promoter inversion (i.e. switched colors). Two dilutions of the cultures were plated (left = lower dilution).

### IR determines the flexibility of flagellar switch in *C. difficile* and impacts *in vivo* virulence

To directly test the ability of the flagellar IR type to alter the virulence of *C. difficile* in mice, we determined the impact of mutating the IR switch region of the reference RT027/ST1 strain, R20291, on its virulence using CRISPR.^39^ To this end, we deleted ∼200 bp of the region upstream of the R20291 *flgB* operon first and then knocked in the wild-type C1 IR (6A/6T), as well as the variants including R2 variants 5A/5T-ON (R2-ON) and 5A/5T-OFF (R2-OFF). It was necessary to generate the R2-ON and R2-OFF variants because our data suggested that the R2 IR sequence greatly reduces the inversion of the flagellar switch region (**Figures 4, 5**). Consistent with our prior data, we observed that the R2-ON resulted in almost all cells in the population carrying the ON orientation (99.9%), while almost all cells in the R2-OFF population carrying the OFF orientation (0.02%) (**Figure S4A**). Furthermore, the R20291_R2-ON (5A/5T-ON) variant exhibited spread using flagellar motility to similar levels as the parental R20291 strain, while the R20291_R2-OFF (5A/5T-OFF) was largely non-motile (**Figure S4B**).

To explore if these R20291 flagellar IR variants have differential virulence in a mouse model, we infected antibiotic-treated mice with the mutant strains and monitored their weight loss (**Figure 6A**). All mutant strains colonized mice to similar levels as the parental R20291 strain, although more variability in colonization levels was observed in strains carrying the flagellar switch in the ON orientation (R20291, **Figure 6B**). Interestingly, fecal CFU levels were below the limit of detection for 4 out of 10 mice infected with R20291_R2-OFF (5A/5T-OFF) and 3 out of 20 mice infected with R20291_R2-ON (5A/5T-ON), implying that the ability to invert the flagellar switch region enhances the initial colonization of *C. difficile* (**Figure 6B**). To focus on the impact of the IR variants on the infection dynamics of *C. difficile*, we excluded these 7 mice from the downstream analyses in mice. All colonized strains maintained high levels of *C. difficile* colonization and shedding up to 14 days post-infection (**Figure 6C and S4C**). Mice infected with the parental R20291 strain exhibited severe weight loss on day 2 post-infection, while the clinical isolate ST1-67, which carries the R2-OFF variant (5A/5T-OFF) did not induce weight loss, consistent with our previous observations(**Figure 6D**).^31^ Mice infected with the “restored” C1 variant, R20291_C1 (6A/6T) phenocopied the weight loss of the parental R20291 strain (**Figure 6D and 6E**), while the R20291 variant carrying R2-OFF, R20291_R2-OFF (5A/5T-OFF), caused minimal weight-loss, essentially phenocopying the avirulence of the ST1-67 strain, which carries the flagellar switch in the R2-OFF orientation (5A/5T-OFF) (**Figure 6D and 6E**).

**Figure 6.**
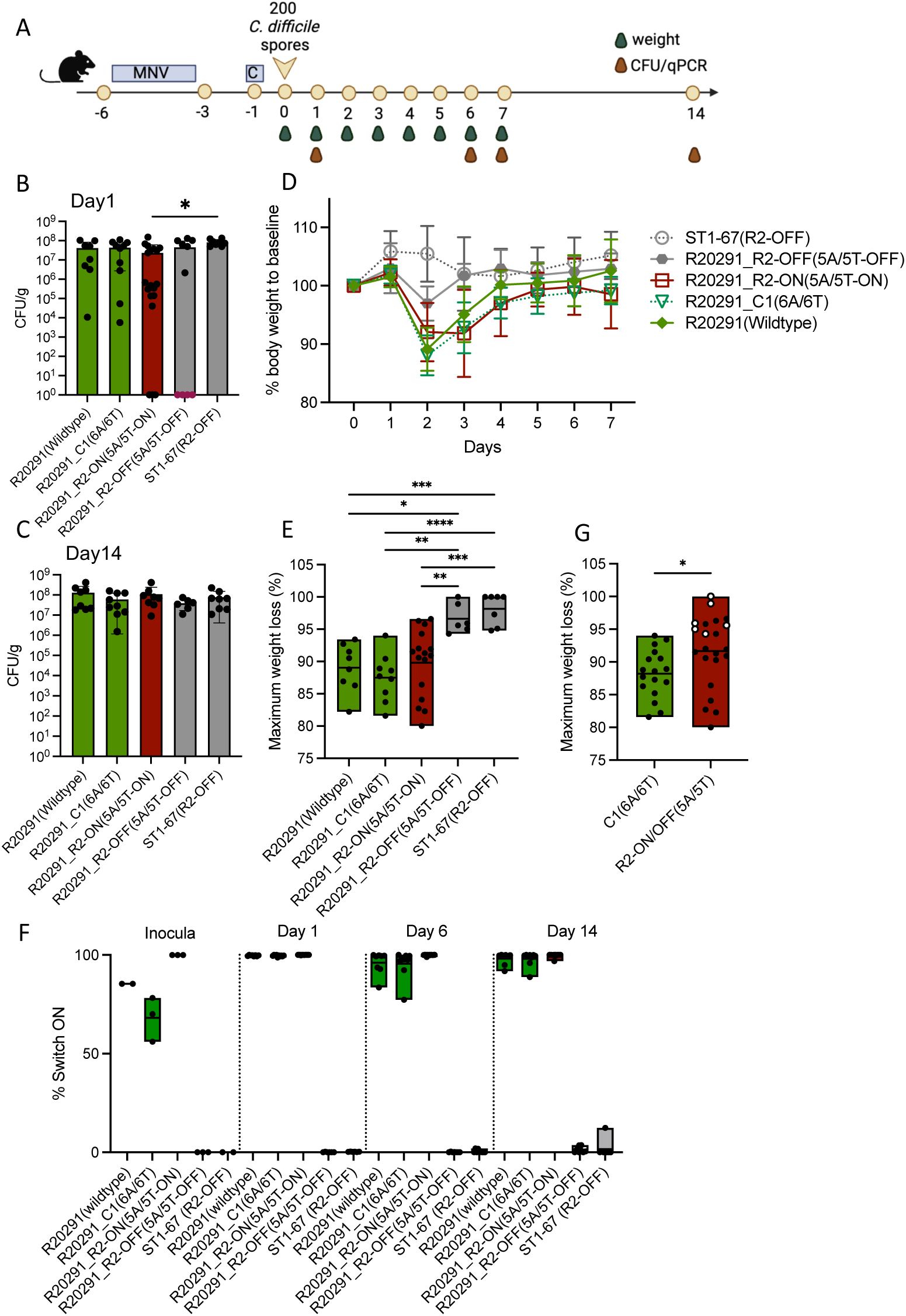
The flagellar repeat variants and flagellar switch orientation impact the virulence of *C. difficile* R20291. (A) Schematic of the experimental procedure. Wild-type C57BL/6 mice (n = 8-20 per group) were treated with Metronidazole, Neomycin and Vancomycin (MNV, 0.25 g/L for each) in drinking water for 3 days, followed by one intraperitoneal injection of Clindamycin (200 mg/mouse), indicated as C in the schematic, 2 days after antibiotic recess. Then, mice were inoculated with 200 indicated *C. difficile* spores via oral gavage. Daily body weight was monitored for 7 days post-infection. (B-C) Fecal colony-forming units were measured by plating on selective agar on 1 day (B) and 14 days (C) post-infection. (D) %Weight loss relative to the baseline of mice infected with the indicated strains up to 7 days post-infection. (E) % Maximum weight loss relative to the baseline of mice infected with indicated strains up to 7 days post-infection. (F) % Flagellar ON vs OFF cells in spore inoculum and in fecal pellets by qPCR. (G) % Maximum weight loss relative to the baseline of mice infected with strains (R20291 background only) grouped by IR types. Statistical significance was calculated by unpaired t-test and One-way ANOVA, * p < 0.05, ** p < 0.01, *** p < 0.001, **** p < 0.0001.

Since orientation-specific qPCR revealed that *C. difficile* strains carrying the R2 IR (5A/5T) IRs remained skewed to either ∼100% ON or ∼100% OFF throughout the infection time course (**Figure 6F**), these data confirm that the R2 IR type largely prevents the flagellar switch from inverting. Notably, since the R20291_R2-OFF variant caused only mild disease, and the only change between the R20291_R2-OFF variant and the parental R20291 strain is a single nucleotide in each of its flagellar switch IR sequences, these data demonstrate that the loss of flagellar gene expression at the population level reduces *C. difficile*’s virulence. In contrast, the R20291_R2-ON (5A/5T-ON) variant exhibited a virulence phenotype in comparison to R20291_R2-OFF (5A/5T-OFF) (**Figure 6D and 6E**), strongly suggesting that flagellar gene expression during infection enhances *C. difficile*’s virulence. However, it should be noted that a majority of mice (11/16, 69%) infected with R20291_R2-ON showed a low-virulence phenotype, causing less than 10% weight loss (which we defined as “low-virulence” for the clinical isolates, **Figure S1**), whereas only 6/17 (35%) of mice infected with the parental R20291 or R20291_C1 variant exhibited less than 10% weight loss. Additionally, pooling the R20291 strains tested by their IR types revealed that *C. difficile* strains carrying the C1 IR type (6A/6T) are significantly more virulent than their counterparts carrying the R2 IR type (5A/5T), regardless of the orientation of the flagellar switch (**Figure 6G**). This is consistent with our finding that clinical isolates carrying the C1 IR type are associated with greater virulence (**Figure 3A**). Taken together, these data demonstrate that the IR types impact the invertibility of *C. difficile* flagellar switch during mouse infection, contributing to heterogeneity in flagellar gene expression at the population level. This heterogeneity appears to promote more severe disease because greatly reducing the invertibility of the flagellar switch region by introducing the R2 IR type variant reduces disease severity in the R20291 strain background, particularly when the R2 IR variant is fixed in the OFF orientation (**Figure 6**).

## DISCUSSION

*C. difficile* infection induces a wide range of disease severity in patients, yet the molecular basis for the variation is poorly understood. Here, we applied a mouse model to study how genetic variation between *C. difficile* isolates impacts their virulence. Using a collection of more than 60 ST1/RT027 clinical isolates, we discovered that variation in the inverted repeat (IR) regions flanking *C. difficile*’s flagellar switch region regulates the invertibility of this switch by the RecV recombinase. Our data indicate that the most common variant found in ST1 strains, C1 (6A/6T), results in greater heterogeneity in flagellar gene expression within a population, with ∼70% of the population carrying the flagellar switch in the ON orientation (**Figures 4, 6**). We further showed that the C1 variant leads to greater disease severity relative to strains with shorter IR variants (**Figure 3, 6G**). The shorter IR variant types - C2 (6A/5T), R1 (5A/6T), and R2 (5A/5T) - restrict the RecV-mediated invertibility of the flagellar switch, effectively “locking” the flagellar switch region into either the ON or OFF orientation (**Figure 4 and 5**). Notably, the decreased invertibility of the flagellar switch region was associated with decreased virulence in ST1 clinical isolates (**Figure 3**) as well as in CRISPR-engineered R20291 strains (**Figure 6**), especially when the engineered R2 switch was in the OFF orientation. Since the IR sequences are sufficient to regulate the inversion of promoter regions in *E. coli* (**Figure 5**) and *C. difficile* R20291 (**Figure 6**), our data reveal that flagellar switch regions IR types control the inversion frequency of this locus (i.e., phase variation), which impacts the virulence of *C. difficile* ST1 strains. In demonstrating that “fixing” the flagellar switch region in the OFF orientation can render ST1 strains avirulent (**Figure 6**), our study reveals another mechanism by which RT027 strains can cause disease of varying severity, which may help explain the wide range of disease outcomes observed in CDI patients.

The invertibility of the flagellar switch likely confers an evolutionary advantage, as more than 95% of isolates worldwide harbor C1 (6A/6T) IR type. Interestingly, in both our strain collection and in the CDC HAI-Seq *C. difficile* strain collection, we found that the C2 (6A/5T)- OFF IR variant accounts for approximately 30% of isolates. This could be due to rigorous testing and *C. difficile* cultivation in the U.S.,^40,41^ as C2-OFF isolates are mostly non-flagellated and are associated with reduced virulence. Since flagellar synthesis and motility is costly^42^ and flagella are targeted by mucosal innate immune response,^19,43^ reducing flagellar motility could be energetically favorable and allow *C. difficile* to evade the immune system as a strategy to increase the persistence of *C. difficile*. However, the ability to switch between flagellar ON and OFF (and vice versa) could enhance *C. difficile*’s fitness during infection by allowing sub-populations of *C. difficile* expressing flagellar genes to inhabit different locations within the gut or function at different stages during the infection process. For example, flagellar motility may help bring *C. difficile* cells closer to the host epithelium to improve toxin binding to its target cells, which could explain why C1 (6A/6T) isolates, which have the most flexible flagellar switch, exhibit the highest virulence among the ST1 strains tested. Future investigations could focus on examining the relative locations of flagellar-ON vs. flagellar-OFF cells for *C. difficile* with various IRs to gain mechanistic insight into the relationship between flagellar heterogeneity and *in vivo* virulence.

Bacteria apply phase variation to reach population phenotypical heterogeneity to promote adaptation to various environmental conditions. Such heterogeneity may increase motility, antibiotic resistance and pathogenic potentials.^44,45^ Notably in *C. difficile*, RecV regulates multiple phase variable regions including flagella as well as a cell wall protein CwpV.^46,47^ CwpV deposited on a proteinaceous layer on the cell surface of *C. difficile* promotes bacterial aggregation *in vitro* and potentially promotes intestinal colonization.^48^ Moreover, *C. difficile* regulates it surface motility via regulation by CmrRST system, which induces cell chaining phenotype on surfaces and contributes to disease development in hamsters.^49,50^ Here, we observed the heterogeneity of *C. difficile* flagella upon infection in mice and the flagellar heterogeneity is important for *C. difficile* virulence. Together, we and others underscore the importance of phenotypical heterogeneity in bacterial survival and virulence and encourage more studies on such phase variable traits of otherwise genetically identical strains.

*C. difficile* flagellar expression has been also linked to toxin production. Turning on the expression of flagellar operon increases the expression of SigD, which is a sigma factor that not only further regulates flagellar synthesis and motility but also positively regulates *tcdR* for toxin production.^22^ Thus, whether flagellar expression contributes to virulence beyond regulating toxin levels is unclear. Here, we demonstrated that the engineered R20291 mutant with a biased ON flagellar orientation is significantly more virulent than its counterpart with a biased OFF flagellar orientation, supporting the role of flagella in *C. difficile* virulence. However, the increase of virulence by tuning ON flagella did not surpass the virulence of *C. difficile* with heterogeneous flagellar orientations, as for both clinical isolates and CRISPR-engineered R20291 mutants. Moreover, we did not find an association between toxin levels in fecal samples and the degree of virulence on a panel of 22 isolates. These suggest that the expression of flagellar (and with coupled toxin expression) provides a relatively minor contribution to virulence and is likely to be strain-dependent, while the invertibility of the flagellar switch may provide major fitness and impact on *C. difficile* virulence (i.e. via impacting the locations of toxin production).

The flagellar phase variation is regulated by DNA recombinase RecV.^21,51^ RecV reversibly inverts the flagellar switch between ON and OFF orientations, and the proportion of ON vs. OFF in the resulting population is likely selected by the residing environments.^52^ Thus, the flexible switch leads to more variable ON and OFF compositions spanning different conditions (from vegetative broth culture and prepared spores to mouse infection), while the less flexible switches result in their highly biased orientation all the time, regardless of the conditions. Using a heterogenous *E. coli* strain, we demonstrated that the flagellar IR type variants reduce RecV-mediated DNA inversion. This is further confirmed with CRIPSR-engineered mutants as R20291 with R2-5A/5T IRs demonstrated highly biased flagellar orientations. RecV targets IRs for inversion, and the potential recombination sites were speculated to be at the borders of these IRs.^51^ Notably, deleting the 1^st^ T and its adjacent nucleotides (Δ3) also “locked” the flagellar orientation into ON or OFF.^23^ Together with our observation, lacking one T in the 6T track in Right IR is likely impacting the efficiency of RecV function. Further experiments should dissect out at what steps that the IR variants impact RecV, such as sequence-specific DNA binding, DNA cleavage, strand exchange, or religation.

In summary, our study identifies the flagellar switch IR as a determinant of the heterogeneity *C. difficile* flagellar phase variation, which also introduces variable virulence outcomes within a single RT027 strain type. Since the invertibility of the flagellar switch is highly associated with the virulence of clinical isolates, we highlight the potential of using flagellar switch inverted repeats as an easily accessible genetic trait to predict pathogen virulence. Further research on the flagellar switch regions of clinical non-ST1 strains may provide additional insights into the wide range of disease severities in patients infected with *C. difficile*.

## Acknowledgement

We would like to thank Dr. Rita Tamayo for providing the primers and reference strains for the flagellar orientation-specific qPCR. We thank Dr. Craig D. Ellermeier for generously providing us the CRISPR editing system of *C. difficile* and Dr. Louis-Charles Fortier for providing *C. difficile* R20291 strain. We thank Dr. Femi Olorunniji for providing the recombinase-switch plasmid. We thank the animal facilities of the University of Chicago and Tufts University for their help with mouse experiments. We thank the Pamer lab members and Shen lab members for helpful discussions. This work was supported by National Institutes of Health R01 AI095706 (to E.G.P), R21 AI168849 (to A.S), R35 GM149586 (to P.A.R), the Duchossois Family Institute of the University of Chicago, and a Burroughs Wellcome Fund Investigators in the Pathogenesis of Disease Award to A.S. The funders had no role in study design, data collection, and interpretation, or the decision to submit the work for publication. The graphical schematics were created with BioRender. com.

## Author contributions

N.T.Q.N, Q.D., A.S. and E.G.P. conceived the project. N.T.Q.N, Q.D., H.L. and P.A.R. analyzed the data. N.T.Q.N., Q.D., Y.P., J.K.S., and P.K., performed experiments. M.K. isolated *C. difficile* isolates. V.B.Y. and E.S.S. sequenced clinical isolates. N.T.Q.N., Q.D., A.S., P.A.R. and E.G.P. interpreted the results and wrote the manuscript.

## Declaration of generative AI and AI-assisted technologies in the writing process

During the preparation of this work, the author(s) used Grammarly in order to proofread the manuscript. After using this tool/service, the author(s) reviewed and edited the content as needed and take(s) full responsibility for the content of the publication.

## Methods

### Clinical *C. difficile* isolates collection

Clinical *C. difficile* isolates were collected from 2013 to 2017 at Memorial Sloan Kettering Cancer Center (MSKCC) from patients receiving bone marrow transplants and cancer chemotherapy.^40^ The isolates were sequenced whole genome in an Illumina Hiseq platform^33^ and circularized at Duchossois Family Institute by MinION Nanopore sequencing (Oxford Nanopore Technologies). Whole genome sequences are available at National Center for Biotechnology Information, BioProject PRJNA595724.

### Bacterial growth and spore collection

Frozen stock of *C. difficile* was struck onto Brain heart infusion (BHI) agar plus 0.1% (w/v) sodium taurocholate hydrate (Cat. 86339, Sigma-Aldrich). A single colony was picked and sub-cultured in BHI medium (BD, 237500) with yeast extract and 0.1% L-Cysteine (BHIS) at 37°C in anaerobic chamber (Coy lab). To prepare spores for the mouse infections, *C. difficile* was either incubated in BHIS broth for approximately two months or on 70:30 agar for 4-5 days to encourage sporulation.^53^ Next, spores were separated from cell debris by gradient centrifugation using a 20%/50% (wt/vol) HistoDenz (D2158, Sigma-Aldrich) or 50% (wt/vol) sucrose gradient, then washed five times in sterile water at 14,000 x g for 5 minutes.^53^ The spore solution was further incubated at 65°C for 20 minutes to kill vegetative cells. Spore purity was confirmed by the absence of vegetative cell growth on BHIS plates or by microscopy.

### Mouse experiment

C57BL/6 female mice from six- to eight-week-old were purchased from Jackson laboratory and housed in the specific-pathogen-free (SPF) facility at University of Chicago or the Tufts University School of Medicine. Mice were randomized and administered *ad libitum* with a cocktail of metronidazole (0.25g/l), neomycin (0.25g/l), and vancomycin (0.25g/l) for three days. Two days after antibiotics removal, mice were intraperitoneally injected with 200µg/mouse clindamycin. After 24 hours, mice were oral gavage with approximately 200 spores of *C. difficile*. All mice infected with *C. difficile* were single-housed then monitored for weight loss and clinical sign of disease for seven days. CFU/g feces counting, and toxin titer measurement was done from fecal pellet collected one day after infection as described elsewhere.^31^

### RNA sequencing

Mice were infected with spores from the four selected isolates (ST1-6, ST1-12, ST1-27, and ST1-53). After 24 hours, mice were sacrificed to collect cecal content for RNA extraction (Qiagen RNeasy PowerMicrobiome kit). Total RNA was sent to Genewiz for rRNA removal and library preparation before sequencing. Adapters were trimmed off from the raw reads, and their quality was assessed and controlled using Trimmomatic (v.0.39),^54^ then human genome was identified and removed by kneaddata (v0.7.10, https://github.com/biobakery/kneaddata), while ribosomal RNA was removed by aligned the clean non-host reads to silva database (138.1 SSURef).^55–57^ The remaining reads from each sample were mapped to their corresponding circularized genome using bowtie2,^58^ and reads counts of each gene were obtained by running featureCounts from Subread (v2.0.1),^59^ and the core gene counts were normalized by DESeq2.^60^

### Sequencing the inverted-repeat sequences

64 isolates with the flagellar switch sequence resembling that of *C. difficile* R20291 were submitted for Sanger sequencing. Firstly, the ON and OFF sequence of each isolate was amplified using the primer pairs as designed by.^21^ The PCR product was purified using QIAquick PCR purification kit (Cat. 28104, Qiagen). Purified products were sent to University of Chicago DNA Sequencing facility and sequenced using the same primer pairs for PCR reaction.

### Swim plates

*C. difficile* was grown in BHIS broth to late-log phase then diluted to OD_600nm_ ∼0.5. To ensure a small number of *C. difficile* cells was added for testing, we submerged a 10 µL pipette tip into diluted *C. difficile* culture without drawing any liquid. The pipette tip carrying *C. difficile* cells on it was then stabbed into 0.3% agarose buffered with BHIS, pH 7.0. The swim plates were then incubated at 37°C. After 24 hours, the swim zone size was measured and compared between isolates.

### Growth curve

To ensure that only live cells were used for growth curve, we prepared a fresh culture from overnight culture at the ratio 1:100. When the back-diluted culture reached log phase, we subculture it into fresh BHI broth at a 1:100 dilution. The culture was then loaded into a 96-well plate for OD_600_ measurements. The incubation time is 24 hours with OD measurements every 10 minutes at 37^°^C with shaking.

### Transmission electron microscope (TEM)

*C. difficile* was inoculated from frozen stocks onto a BHIS agar plate and incubated overnight. To collect cells for TEM, 10µl of distilled water was dropped onto a colony. After two minutes, a copper grid (CF400-Cu, EMS) was placed on top of the soaked colony so that vegetative cells were passively transferred to the grid. To stain the cells, one drop of uranyl acetate 1% was added to the grid and incubate for 30 seconds. TEM pictures of the stained cells were then taken at 2900X1.4 or 5900X1.4 using FEI Tecnai F30 microscope at University of Chicago Advanced Electron Microscopy Facility.

### Quantitative PCR of the flagellar switch

*C. difficile* was grown in BHIS broth to late log phase (OD_600nm_ ∼ 0.5) with three to four replicates per isolate. *C. difficile* DNA was extracted by Qiamp PowerFecal Pro DNA kit (Cat. 51804). Fecal samples were harvested from infected mice and fecal DNA was extracted either using QIAamp PowerFecal Pro DNA Kit or as previously described.^23^ Quantitative real-time PCR was done using 20-30 ng template and flagellar switch primers.^21^ *C. difficile* R20291 mutated strains with flagellar switch locked ON and OFF were used as the control.^23^ Adenosine kinase (*adk)* gene is used as a reference gene. qPCR was set up using PowerUp™ SYBR™ Green Master Mix (Cat. A25742) and run by QuantStudio Real-Time PCR Systems or NEB Luna qPCR Master Mix and run by Applied Biosystems StepOnePlus system. The switch direction percentage was calculated using the ΔΔCt method as previously reported,^23^ while integrating the primers’ amplification efficiencies.

### Generation of *C. difficile* mutants using CRISPR

CRISPR editing on *C. difficile* strains R20291 was performed as described previously.^39^ Briefly, donor regions for homology were generated by separately amplifying regions ∼500 bp upstream and ∼500 bp downstream of the target of interest. The resulting regions were cloned into pCE677 between NotI and XhoI sites by Gibson Assembly. Geneious Prime (v11) was used to design sgRNAs targeting each deleted target. sgRNA fragments were then amplified by PCR from pCE677, using an upstream primer that introduces the altered guide and inserted at the MscI and MluI sites of the pCE677-derivative with the appropriate homology region. The regions of plasmids constructed using PCR were verified by Sanger sequencing. Plasmids were then passaged through NEBturbo *E. coli* strain before transformation into *Bacillus subtilis* strain BS49. The CRISPR-Cas9 deletion plasmids which harbor the *oriT* (*Tn916*) origin of transfer, were then introduced into *C. difficile* strains by conjugation.^61^ *C. difficile* colonies were then screened for proper mutations in the genomes by PCR and Sanger sequencing. To generate *C. difficile* IR repeats mutants, two rounds of CRISPR editing were conducted. The first round was to delete ∼200 bp region containing the flagellar switch and IRs while introducing an *RFP* landing pad (GGCGCCCAGACCGCTAAACTGAAAGTT) into the place. The second round gRNA targeted the *RFP* landing pad, with the mutant IR variant flagellar switch region (either in the ON or OFF orientation) template supplemented for repair. Primers used for CRISPR editing were included in Table S2.

### Flagella switch alignment and IR type survey in public databases

*C. difficile* isolates (N=131) from BioProject PRJNA595724 and PRJEB2318 (N=1198) were downloaded from NCBI and assembled into contigs using SPAdes.^62^ Eleven of those isolates did not pass the assembling process. In addition, a collection of 66 *C. difficile* genomes from Patric (date: Mar. 15 2022, https://www.bv-brc.org/ were also downloaded. MLST was determined on those contigs by mlst (Seemann T, mlst Github https://github.com/tseemann/mlst).^63^ Flagellar switch region and 50 bp upstream and downstream of 68 isolates in our collection of both the ON and OFF sequences were used as query to BLAST^64^ against the assembled contigs, and hits with at least 85% identity and 85% coverage of the query are considered a valid match.

### *E. coli* colorimetric assay

Plasmids for the E. coli experiments (see Table S3) were ordered from Twist Bioscience and were based on backbone vectors kindly onboarded with Twist by Dr. Femi Olorunniji (Liverpool John Moores University). Plasmids were checked by full-plasmid sequencing (Plasmidsaurus). pQD1 is a pBAD derivative for tightly controlled arabinose-inducible expression of RecV. Test plasmids pQD2-7 are based on pϕC31-invPB described previously,^38^ with IRs from the flagellar switch replacing the att site for ϕC31 integrase.

Assays were performed as described previously with minor variations.^65^ *E. coli* DS941^66^ was co-transformed with pQD1 plus one of the test plasmids, then after recovery grown overnight at 37C in 10ml LB supplemented with 0.2% glucose (to enhance repression of RecV), kanamycin (50 µg/ml; to maintain the test plasmid) and chloramphenicol (30 µg/ml; to maintain pQD1). In the morning, the OD_600_ was ∼2. 200 μl of that culture was diluted into 10ml LB plus 0.2% glucose, kanamycin (50 µg/ml) and chloramphenicol (30 µg/ml) and grown at 37C until the OD_600_ reached ∼0.5. They were then switched to arabinose to induce RecV expression by pelleting the cells, removing the supernatant, and resuspending in 10ml LB plus 0.2% arabinose, kanamycin (50 µg/ml) and chloramphenicol (30 µg/ml), and growing at 37C for 4 hours. When cultures were plated immediately after the RecV induction period the colonies for experiments with pQD1 plus test plasmids containing the 6A/6T IRs (pQD2 or pQD3) were mostly yellow due to mixed populations of substrate and product plasmid within each founder cell (the test plasmid replicates to high copy number within each host cell).

To be separated from one another the test plasmids needed to be recovered then retransformed. After the 4 hours of RecV expression in arabinose, the cultures were switched back to glucose for overnight growth: 1 ml was removed, pelleted, and resuspended in 1 ml LB supplemented with 0.2% glucose and kanamycin (50 µg/ml), then 50ul of that was used to inoculate 5 ml of LB supplemented with 0.2% glucose and kanamycin (50 µg/ml), which was grown overnight. Plasmids were recovered by miniprep. 1ul of each plasmid was used to transform competent DS941 E. coli, then two different volumes were plated on LB plus kanamycin. Colony color was visualized using a ChemiDoc imager (BioRad). The red and green channel images are overlaid in Figure 5.

Full-plasmid sequencing (Plasmidsaurus) was used to verify the recombination products. Plasmids recovered from a green colony resulting from the pQD1 + pQD2 (6A/6T – OFF /red) experiment were identical to pQD3 (6A/6T – ON /green), confirming the expected inversion. To isolate products from the experiments using pQD4 (5A/6T – OFF /red) and pQD5 (5A/6T – ON /green), the 1 or 2 product-color colonies seen in Figure 5 were picked, then restreaked to ensure separation from their substrate-containing neighbors. Sequencing confirmed that product plasmids from the pQD4 experiment matched the sequence of pQD5, and vice versa.

### RNA extraction, reverse transcription and RT-qPCR

Cecal RNA was extracted using Rneasy PowerMicrobiome Kit (Qiagen) according to the manufacturer’s instructions. Complementary DNA was generated using the QuantiTect reverse transcriptase kit (Qiagen) according to the manufacturer’s instructions. Quantitative PCR was performed on complementary DNA using primers with PowerTrack SYBR Green Master Mix (Thermo Fisher). Reactions were run on a QuantStudio 6 pro (Thermo Fisher). Relative abundance was normalized by ΔΔCt. TcdA_qFor 5’-GTATGGATAGGTGGAGAAGTCA-3’; TcdA_qRev 5’-CTCTTCCTCTAGTAGCTGTAATGC-3’^67^; TcdB_qFor 5’- AGCAGTTGAATATAGTGGTTTAGTTAGAGTTG-3’; TcdB_qRev 5’- CATGCTTTTTTAGTTTCTGGATTGAA-3’^68^; FliS1_qFor 5’- TGCAGGACAATGGGCAAAGG-3’; FliS1_qRev 5’- CAGGCAACACATTATCTATTACCTGG-3’; FliE_qFor 5’- AGGCGAAGATGTTTCTATGCA-3’; ACCTTATTCATTTCTTGATATGCATCA-3’ FliE_qRev 5’-

### Statistical analysis

Kruskal-Wallis, T-tests and One-way ANOVA tests was performed to test the difference in maximum weight loss, swim plate, and RT-qPCR results. Statistical significance was determined by using a *P* value of <0.05. Linear regression analysis was used to estimate the correlation between IR type and weight loss.

## REFERENCES

1. CDC, A. (2019). Antibiotic resistance threats in the United States. US Department of Health and Human Services: Washington, DC, USA.

2. CDC (2020). 2020 Annual Report.

3. Abt, M.C., McKenney, P.T., and Pamer, E.G. (2016). *Clostridium difficile* colitis: pathogenesis and host defence. Nat Rev Microbiol 14, 609–620. 10.1038/nrmicro.2016.108.

4. Lyerly, D.M., Krivan, H.C., and Wilkins, T.D. (1988). *Clostridium difficile*: its disease and toxins. Clin Microbiol Rev 1, 1–18.

5. Ciesielski-Treska, J., Ulrich, G., Rihn, B., and Aunis, D. (1989). Mechanism of action of *Clostridium difficile* toxin B: role of external medium and cytoskeletal organization in intoxicated cells. Eur J Cell Biol 48, 191–202.

6. Florin, I., and Thelestam, M. (1983). Internalization of *Clostridium difficile* cytotoxin into cultured human lung fibroblasts. Biochim Biophys Acta 763, 383–392. 10.1016/0167-4889(83)90100-3.

7. Henriques, B., Florin, I., and Thelestam, M. (1987). Cellular internalisation of *Clostridium difficile* toxin A. Microb Pathog 2, 455–463. 10.1016/0882-4010(87)90052-0.

8. Just, I., Selzer, J., Hofmann, F., Green, G.A., and Aktories, K. (1996). Inactivation of Ras by *Clostridium sordellii* lethal toxin-catalyzed glucosylation. J Biol Chem 271, 10149–10153. 10.1074/jbc.271.17.10149.

9. Just, I., Selzer, J., Wilm, M., von Eichel-Streiber, C., Mann, M., and Aktories, K. (1995). Glucosylation of Rho proteins by *Clostridium difficile* toxin B. Nature 375, 500–503. 10.1038/375500a0.

10. Just, I., Wilm, M., Selzer, J., Rex, G., von Eichel-Streiber, C., Mann, M., and Aktories, K. (1995). The enterotoxin from *Clostridium difficile* (ToxA) monoglucosylates the Rho proteins. J Biol Chem 270, 13932–13936. 10.1074/jbc.270.23.13932.

11. Just, I., Fritz, G., Aktories, K., Giry, M., Popoff, M.R., Boquet, P., Hegenbarth, S., and von Eichel-Streiber, C. (1994). *Clostridium difficile* toxin B acts on the GTP-binding protein Rho. J Biol Chem 269, 10706–10712.

12. Schnizlein, M.K., and Young, V.B. (2022). Capturing the environment of the *Clostridioides difficile* infection cycle. Nat Rev Gastroenterol Hepatol 19, 508–520. 10.1038/s41575-022-00610-0.

13. Vedantam, G., Clark, A., Chu, M., McQuade, R., Mallozzi, M., and Viswanathan, V.K. (2012). *Clostridium difficile* infection: toxins and non-toxin virulence factors, and their contributions to disease establishment and host response. Gut Microbes 3, 121–134. 10.4161/gmic.19399.

14. Burdon, D.W., George, R.H., Mogg, G.A., Arabi, Y., Thompson, H., Johnson, M., Alexander-Williams, J., and Keighley, M.R. (1981). Faecal toxin and severity of antibiotic-associated pseudomembranous colitis. J Clin Pathol 34, 548–551. 10.1136/jcp.34.5.548.

15. La Ragione, R.M., Sayers, A.R., and Woodward, M.J. (2000). The role of fimbriae and flagella in the colonization, invasion and persistence of *Escherichia coli* O78:K80 in the day-old-chick model. Epidemiol Infect 124, 351–363. 10.1017/s0950268899004045.

16. Eaves-Pyles, T., Murthy, K., Liaudet, L., Virag, L., Ross, G., Soriano, F.G., Szabo, C., and Salzman, A.L. (2001). Flagellin, a novel mediator of *Salmonella*-induced epithelial activation and systemic inflammation: I kappa B alpha degradation, induction of nitric oxide synthase, induction of proinflammatory mediators, and cardiovascular dysfunction. J Immunol 166, 1248–1260. 10.4049/jimmunol.166.2.1248.

17. Horstmann, J.A., Lunelli, M., Cazzola, H., Heidemann, J., Kuhne, C., Steffen, P., Szefs, S., Rossi, C., Lokareddy, R.K., Wang, C., et al. (2020). Methylation of *Salmonella* Typhimurium flagella promotes bacterial adhesion and host cell invasion. Nat Commun 11, 2013. 10.1038/s41467-020-15738-3.

18. Baban, S.T., Kuehne, S.A., Barketi-Klai, A., Cartman, S.T., Kelly, M.L., Hardie, K.R., Kansau, I., Collignon, A., and Minton, N.P. (2013). The Role of Flagella in *Clostridium difficile* Pathogenesis: Comparison between a Non-Epidemic and an Epidemic Strain. PLoS One 8, e73026. 10.1371/journal.pone.0073026.

19. Batah, J., Kobeissy, H., Bui Pham, P.T., Denève-Larrazet, C., Kuehne, S., Collignon, A., Janoir-Jouveshomme, C., Marvaud, J.-C., and Kansau, I. (2017). *Clostridium difficile* flagella induce a pro-inflammatory response in intestinal epithelium of mice in cooperation with toxins. Sci Rep 7, 3256. 10.1038/s41598-017-03621-z.

20. Dingle, T.C., Mulvey, G.L., and Armstrong, G.D. (2011). Mutagenic Analysis of the *Clostridium difficile* Flagellar Proteins, FliC and FliD, and Their Contribution to Virulence in Hamsters ▿. Infect Immun 79, 4061–4067. 10.1128/IAI.05305-11.

21. Anjuwon-Foster, B.R., and Tamayo, R. (2017). A genetic switch controls the production of flagella and toxins in *Clostridium difficile*. PLOS Genetics 13, e1006701. 10.1371/journal.pgen.1006701.

22. El Meouche, I., Peltier, J., Monot, M., Soutourina, O., Pestel-Caron, M., Dupuy, B., and Pons, J.-L. (2013). Characterization of the SigD Regulon of *C. difficile* and Its Positive Control of Toxin Production through the Regulation of tcdR. PLoS One 8, e83748. 10.1371/journal.pone.0083748.

23. Trzilova, D., Warren, M.A.H., Gadda, N.C., Williams, C.L., and Tamayo, R. Flagellum and toxin phase variation impacts intestinal colonization and disease development in a mouse model of *Clostridioides difficile* infection. Gut Microbes 14, 2038854. 10.1080/19490976.2022.2038854.

24. Tenover, F.C., Tickler, I.A., and Persing, D.H. (2012). Antimicrobial-resistant strains of *Clostridium difficile* from North America. Antimicrob Agents Chemother 56, 2929–2932. 10.1128/AAC.00220-12.

25. Wieczorkiewicz, J.T., Lopansri, B.K., Cheknis, A., Osmolski, J.R., Hecht, D.W., Gerding, D.N., and Johnson, S. (2016). Fluoroquinolone and Macrolide Exposure Predict *Clostridium difficile* Infection with the Highly Fluoroquinolone- and Macrolide-Resistant Epidemic *C. difficile* Strain BI/NAP1/027. Antimicrob Agents Chemother 60, 418–423. 10.1128/AAC.01820-15.

26. Hubert, B., Loo, V.G., Bourgault, A.M., Poirier, L., Dascal, A., Fortin, E., Dionne, M., and Lorange, M. (2007). A portrait of the geographic dissemination of the *Clostridium difficile* North American pulsed-field type 1 strain and the epidemiology of *C. difficile*-associated disease in Quebec. Clin Infect Dis 44, 238–244. 10.1086/510391.

27. Labbe, A.C., Poirier, L., Maccannell, D., Louie, T., Savoie, M., Beliveau, C., Laverdiere, M., and Pepin, J. (2008). *Clostridium difficile* infections in a Canadian tertiary care hospital before and during a regional epidemic associated with the BI/NAP1/027 strain. Antimicrob Agents Chemother 52, 3180–3187. 10.1128/AAC.00146-08.

28. Morgan, O.W., Rodrigues, B., Elston, T., Verlander, N.Q., Brown, D.F., Brazier, J., and Reacher, M. (2008). Clinical severity of *Clostridium difficil*e PCR ribotype 027: a case-case study. PLoS One 3, e1812. 10.1371/journal.pone.0001812.

29. Sirard, S., Valiquette, L., and Fortier, L.C. (2011). Lack of association between clinical outcome of *Clostridium difficile* infections, strain type, and virulence-associated phenotypes. J Clin Microbiol 49, 4040–4046. 10.1128/JCM.05053-11.

30. Carlson, P.E., Walk, S.T., Bourgis, A.E., Liu, M.W., Kopliku, F., Lo, E., Young, V.B., Aronoff, D.M., and Hanna, P.C. (2013). The relationship between phenotype, ribotype, and clinical disease in human *Clostridium difficile* isolates. Anaerobe 24, 109–116. 10.1016/j.anaerobe.2013.04.003.

31. Dong, Q., Lin, H., Allen, M.-M., Garneau, J.R., Sia, J.K., Smith, R.C., Haro, F., McMillen, T., Pope, R.L., Metcalfe, C., et al. (2023). Virulence and genomic diversity among clinical isolates of ST1 (BI/NAP1/027) *Clostridioides difficile*. Cell Rep 42, 112861. 10.1016/j.celrep.2023.112861.

32. Paulick, A., Adamczyk, M., Anderson, K., Vlachos, N., Machado, M.J., McAllister, G., Korhonen, L., Guh, A.Y., Halpin, A.L., Rasheed, J.K., et al. (2021). Characterization of *Clostridioides difficile* Isolates Available through the CDC & FDA Antibiotic Resistance Isolate Bank. Microbiol Resour Announc 10. 10.1128/MRA.01011-20.

33. Miles-Jay, A., Young, V.B., Pamer, E.G., Savidge, T.C., Kamboj, M., Garey, K.W., and Snitkin, E.S. (2021). A multisite genomic epidemiology study of *Clostridioides difficile* infections in the USA supports differential roles of healthcare versus community spread for two common strains. Microb Genom 7, 000590. 10.1099/mgen.0.000590.

34. Dingle, K.E., Didelot, X., Quan, T.P., Eyre, D.W., Stoesser, N., Golubchik, T., Harding, R.M., Wilson, D.J., Griffiths, D., Vaughan, A., et al. (2017). Effects of control interventions on *Clostridium difficile* infection in England: an observational study. Lancet Infect Dis 17, 411–421. 10.1016/S1473-3099(16)30514-X.

35. Eyre, D.W., Davies, K.A., Davis, G., Fawley, W.N., Dingle, K.E., De Maio, N., Karas, A., Crook, D.W., Peto, T.E.A., Walker, A.S., et al. (2018). Two Distinct Patterns of *Clostridium difficile* Diversity Across Europe Indicating Contrasting Routes of Spread. Clin Infect Dis 67, 1035–1044. 10.1093/cid/ciy252.

36. Eyre, D.W., Didelot, X., Buckley, A.M., Freeman, J., Moura, I.B., Crook, D.W., Peto, T.E.A., Walker, A.S., Wilcox, M.H., and Dingle, K.E. (2019). *Clostridium difficile* trehalose metabolism variants are common and not associated with adverse patient outcomes when variably present in the same lineage. EBioMedicine 43, 347–355. 10.1016/j.ebiom.2019.04.038.

37. Stoesser, N., Crook, D.W., Fung, R., Griffiths, D., Harding, R.M., Kachrimanidou, M., Keshav, S., Peto, T.E., Vaughan, A., Walker, A.S., et al. (2011). Molecular epidemiology of *Clostridium difficile* strains in children compared with that of strains circulating in adults with *Clostridium difficile*-associated infection. J Clin Microbiol 49, 3994–3996. 10.1128/JCM.05349-11.

38. Olorunniji, F.J., Lawson-Williams, M., McPherson, A.L., Paget, J.E., Stark, W.M., and Rosser, S.J. (2019). Control of ϕC31 integrase-mediated site-specific recombination by protein trans-splicing. Nucleic Acids Res 47, 11452–11460. 10.1093/nar/gkz936.

39. Kaus, G.M., Snyder, L.F., Müh, U., Flores, M.J., Popham, D.L., and Ellermeier, C.D. (2020). Lysozyme Resistance in *Clostridioides difficile* Is Dependent on Two Peptidoglycan Deacetylases. J Bacteriol 202, e00421–20. 10.1128/JB.00421-20.

40. Kamboj, M., McMillen, T., Syed, M., Chow, H.Y., Jani, K., Aslam, A., Brite, J., Fanelli, B., Hasan, N.A., Dadlani, M., et al. (2021). Evaluation of a Combined Multilocus Sequence Typing and Whole-Genome Sequencing Two-Step Algorithm for Routine Typing of *Clostridioides difficile*. J Clin Microbiol 59. 10.1128/JCM.01955-20.

41. Tariq, R., Weatherly, R.M., Kammer, P.P., Pardi, D.S., and Khanna, S. (2017). Experience and Outcomes at a Specialized *Clostridium difficile* Clinical Practice. Mayo Clin Proc Innov Qual Outcomes 1, 49–56. 10.1016/j.mayocpiqo.2017.05.002.

42. Schavemaker, P.E., and Lynch, M. (2022). Flagellar energy costs across the tree of life. Elife 11. 10.7554/eLife.77266.

43. Batah, J., Denève-Larrazet, C., Jolivot, P.-A., Kuehne, S., Collignon, A., Marvaud, J.-C., and Kansau, I. (2016). *Clostridium difficile* flagella predominantly activate TLR5-linked NF-κB pathway in epithelial cells. Anaerobe 38, 116–124. 10.1016/j.anaerobe.2016.01.002.

44. Jiang, X., Hall, A.B., Arthur, T.D., Plichta, D.R., Covington, C.T., Poyet, M., Crothers, J., Moses, P.L., Tolonen, A.C., Vlamakis, H., et al. (2019). Invertible promoters mediate bacterial phase variation, antibiotic resistance, and host adaptation in the gut. Science 363, 181–187. 10.1126/science.aau5238.

45. Krinos, C.M., Coyne, M.J., Weinacht, K.G., Tzianabos, A.O., Kasper, D.L., and Comstock, L.E. (2001). Extensive surface diversity of a commensal microorganism by multiple DNA inversions. Nature 414, 555–558. 10.1038/35107092.

46. Emerson, J.E., Reynolds, C.B., Fagan, R.P., Shaw, H.A., Goulding, D., and Fairweather, N.F. (2009). A novel genetic switch controls phase variable expression of CwpV, a *Clostridium difficile* cell wall protein. Mol Microbiol 74, 541–556. 10.1111/j.1365-2958.2009.06812.x.

47. Sekulovic, O., Ospina Bedoya, M., Fivian-Hughes, A.S., Fairweather, N.F., and Fortier, L.-C. (2015). The *Clostridium difficile* cell wall protein CwpV confers phase-variable phage resistance. Mol Microbiol 98, 329–342. 10.1111/mmi.13121.

48. Reynolds, C.B., Emerson, J.E., de la Riva, L., Fagan, R.P., and Fairweather, N.F. (2011). The *Clostridium difficil*e cell wall protein CwpV is antigenically variable between strains, but exhibits conserved aggregation-promoting function. PLoS Pathog 7, e1002024. 10.1371/journal.ppat.1002024.

49. Garrett, E.M., Mehra, A., Sekulovic, O., and Tamayo, R. (2021). Multiple Regulatory Mechanisms Control the Production of CmrRST, an Atypical Signal Transduction System in *Clostridioides difficile*. mBio 13, e0296921. 10.1128/mbio.02969-21.

50. Ribis, J.W., Nieto-Acuna, C.A., DiBenedetto, N., Mehra, A., Dong, Q., Nagawa, I.L., Meouche, I.E., Aldridge, B.B., Dunlop, M.J., Tamayo, R., et al. (2024). Unique growth and morphology properties of Clade 5 *Clostridioides difficile* strains revealed by single-cell time-lapse microscopy. Preprint at bioRxiv, 10.1101/2024.02.13.580212 10.1101/2024.02.13.580212.

51. Sekulovic, O., Garrett, E.M., Bourgeois, J., Tamayo, R., Shen, A., and Camilli, A. (2018). Genome-wide detection of conservative site-specific recombination in bacteria. PLOS Genetics 14, e1007332. 10.1371/journal.pgen.1007332.

52. Trzilova, D., and Tamayo, R. (2021). Site-specific recombination – how simple DNA inversions produce complex phenotypic heterogeneity in bacterial populations. Trends Genet 37, 59–72. 10.1016/j.tig.2020.09.004.

53. Weldy, M., Evert, C., Dosa, P.I., Khoruts, A., and Sadowsky, M.J. (2020). Convenient Protocol for Production and Purification of *Clostridioides difficile* Spores for Germination Studies. STAR Protoc 1, 100071. 10.1016/j.xpro.2020.100071.

54. Bolger, A.M., Lohse, M., and Usadel, B. (2014). Trimmomatic: a flexible trimmer for Illumina sequence data. Bioinformatics 30, 2114–2120. 10.1093/bioinformatics/btu170.

55. Quast, C., Pruesse, E., Yilmaz, P., Gerken, J., Schweer, T., Yarza, P., Peplies, J., and Glöckner, F.O. (2013). The SILVA ribosomal RNA gene database project: improved data processing and web-based tools. Nucleic Acids Research 41, D590–D596. 10.1093/nar/gks1219.

56. Glöckner, F.O., Yilmaz, P., Quast, C., Gerken, J., Beccati, A., Ciuprina, A., Bruns, G., Yarza, P., Peplies, J., Westram, R., et al. (2017). 25 years of serving the community with ribosomal RNA gene reference databases and tools. Journal of Biotechnology 261, 169–176. 10.1016/j.jbiotec.2017.06.1198.

57. Yilmaz, P., Parfrey, L.W., Yarza, P., Gerken, J., Pruesse, E., Quast, C., Schweer, T., Peplies, J., Ludwig, W., and Glöckner, F.O. (2014). The SILVA and “All-species Living Tree Project (LTP)” taxonomic frameworks. Nucleic Acids Research 42, D643–D648. 10.1093/nar/gkt1209.

58. Langmead, B., and Salzberg, S.L. (2012). Fast gapped-read alignment with Bowtie 2. Nat Methods 9, 357–359. 10.1038/nmeth.1923.

59. Liao, Y., Smyth, G.K., and Shi, W. (2014). featureCounts: an efficient general purpose program for assigning sequence reads to genomic features. Bioinformatics 30, 923–930. 10.1093/bioinformatics/btt656.

60. Love, M.I., Huber, W., and Anders, S. (2014). Moderated estimation of fold change and dispersion for RNA-seq data with DESeq2. Genome Biol 15, 550. 10.1186/s13059-014-0550-8.

61. McAllister, K.N., Bouillaut, L., Kahn, J.N., Self, W.T., and Sorg, J.A. (2017). Using CRISPR-Cas9-mediated genome editing to generate *C. difficile* mutants defective in selenoproteins synthesis. Sci Rep 7, 14672. 10.1038/s41598-017-15236-5.

62. Prjibelski, A., Antipov, D., Meleshko, D., Lapidus, A., and Korobeynikov, A. (2020). Using SPAdes De Novo Assembler. Curr Protoc Bioinformatics 70, e102. 10.1002/cpbi.102.

63. Griffiths, D., Fawley, W., Kachrimanidou, M., Bowden, R., Crook, D.W., Fung, R., Golubchik, T., Harding, R.M., Jeffery, K.J.M., Jolley, K.A., et al. (2010). Multilocus sequence typing of *Clostridium difficile*. J Clin Microbiol 48, 770–778. 10.1128/JCM.01796-09.

64. Camacho, C., Coulouris, G., Avagyan, V., Ma, N., Papadopoulos, J., Bealer, K., and Madden, T.L. (2009). BLAST+: architecture and applications. BMC Bioinformatics 10, 421. 10.1186/1471-2105-10-421.

65. Shin, H., Holland, A., Alsaleh, A., Retiz, A.D., Pigli, Y.Z., Taiwo-Aiyerin, O.T., Reyes, T.P., Bello, A.J., Olorunniji, F.J., and Rice, P.A. (2024). Identification of cognate recombination directionality factors for large serine recombinases by virtual pulldown. Preprint at bioRxiv, 10.1101/2024.06.11.598349 10.1101/2024.06.11.598349.

66. Summers, D.K., and Sherratt, D.J. (1988). Resolution of ColE1 dimers requires a DNA sequence implicated in the three-dimensional organization of the cer site. EMBO J 7, 851– 858. 10.1002/j.1460-2075.1988.tb02884.x.

67. Babakhani, F., Bouillaut, L., Sears, P., Sims, C., Gomez, A., and Sonenshein, A.L. (2013). Fidaxomicin inhibits toxin production in *Clostridium difficile*. Journal of Antimicrobial Chemotherapy 68, 515–522. 10.1093/jac/dks450.

68. Wroblewski, D., Hannett, G.E., Bopp, D.J., Dumyati, G.K., Halse, T.A., Dumas, N.B., and Musser, K.A. (2009). Rapid molecular characterization of *Clostridium difficile* and assessment of populations of *C. difficile* in stool specimens. J Clin Microbiol 47, 2142–2148. 10.1128/JCM.02498-08.

