## Supplemental figures S1-S4 and Table S1-S3 for "Flagellar switch inverted repeat impacts flagellar invertibility and varies *Clostridioides difficile* RT027/MLST1 virulence"

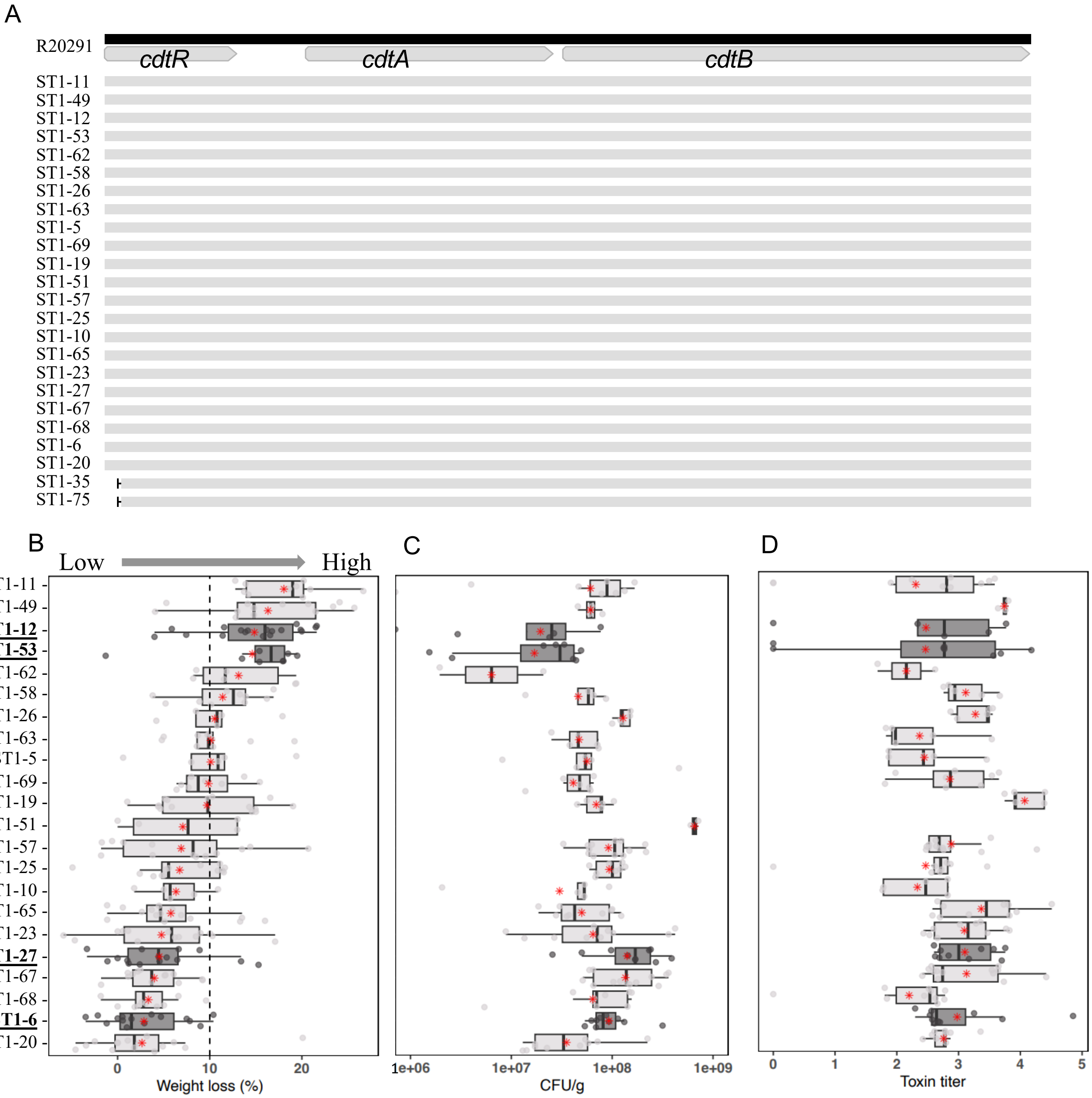


**Figure S1**: **Variable virulence of C. difficile ST1 isolates in antibiotic-treated mice**. (A) *cdtLoc* alignments of the indicated ST1 strains. ST1-35 and ST1-75 carry a 69 bp in-frame deletion of *cdtR*. (B) The maximum weight loss percentage (n = 5 to 13 mice per isolate). In antibiotic-treated mice infected with *C. difficil*e ST1 strains, the most severe weight loss occurs on day two or three post-infection. (C) Boxplot showing the CFU/g data from fecal pellets collected one day after infection (n = 2 to 13 mice per isolate). (D) Toxin titer from fecal pellets collected one day after infection (n = 2 to 13 mice per isolate, except for ST1-51). Each black dot represents for a data point from one mouse. The red asterisk represents for an average value. The dash line in panel B shows the 10% cut-off of weight loss, with strains causing < 10% weight-loss on average being deemed ”low-virulence” strains. ST1-12, ST1-53, ST1-27 and ST1-6 were highlighted in bold and dark-grey plots.

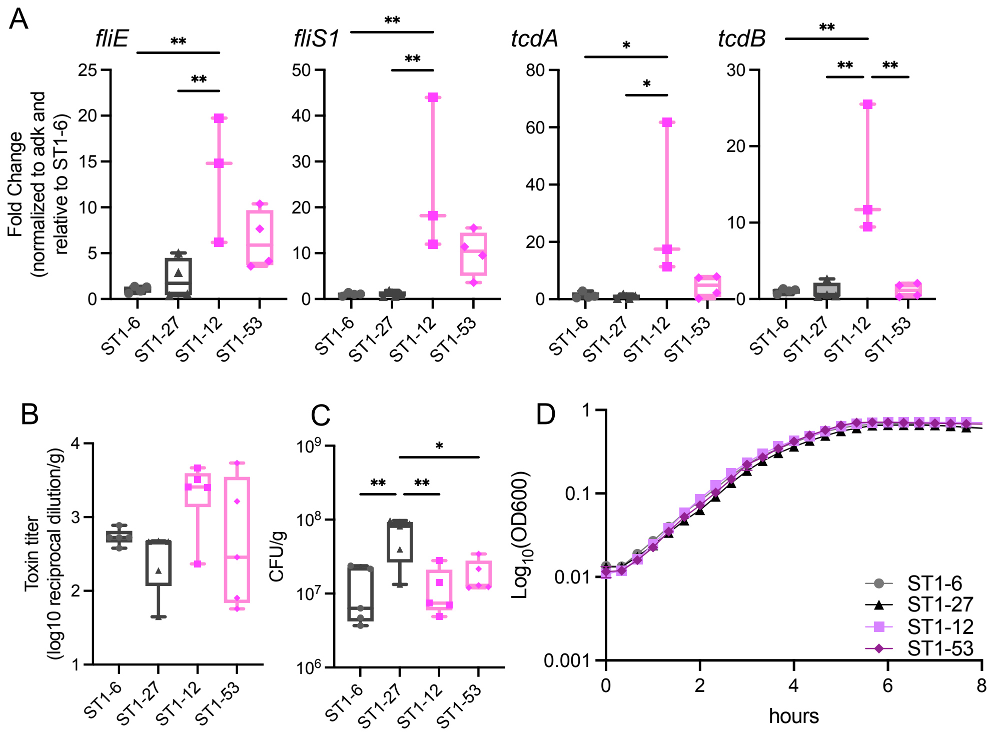


**<u>Figure S2</u>**: High-virulence isolates over-express flagellar genes relative to low-virulence. (A) RT-qPCR confirms the difference in flagellar gene expression between high- (pink) and low-virulence (black) isolates. Fold-change of flagellar genes *fliE*, *fliS1*, and toxin genes *tcdA*, and *tcdB* were normalized to the housekeeping gene, *adk,* relative to ST1_6 (n=3 to 4 RT-qPCR for each isolate). (B) The toxin titer and (C) CFU/g generated from fecal content collected from mice one day post infection (n = 5 mice per isolate), relate to Fig. 1B. (D) Growth curve of 4 isolates in BHIS broth. Statistical significance was calculated by One-way ANOVA, * $p < 0.05$. ** $p < 0.01$.

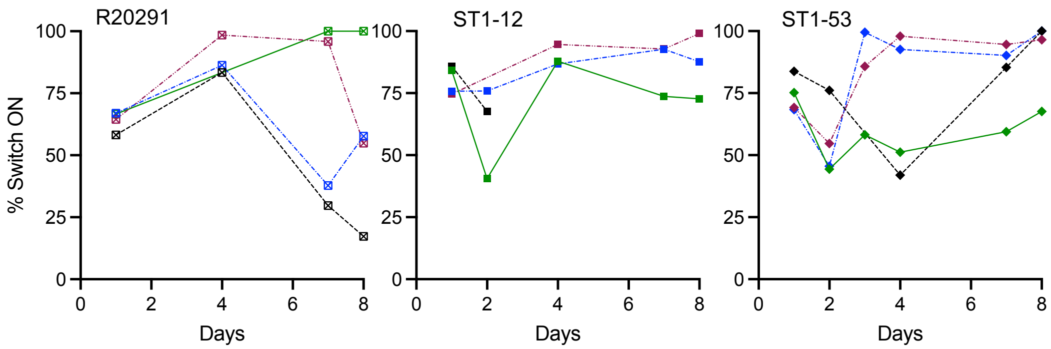


**Figure S3**: Flagellar inverted repeat types are associated with flagellar switch invertibility in mice. C1 data from Figure. 3C shown by individual mouse.

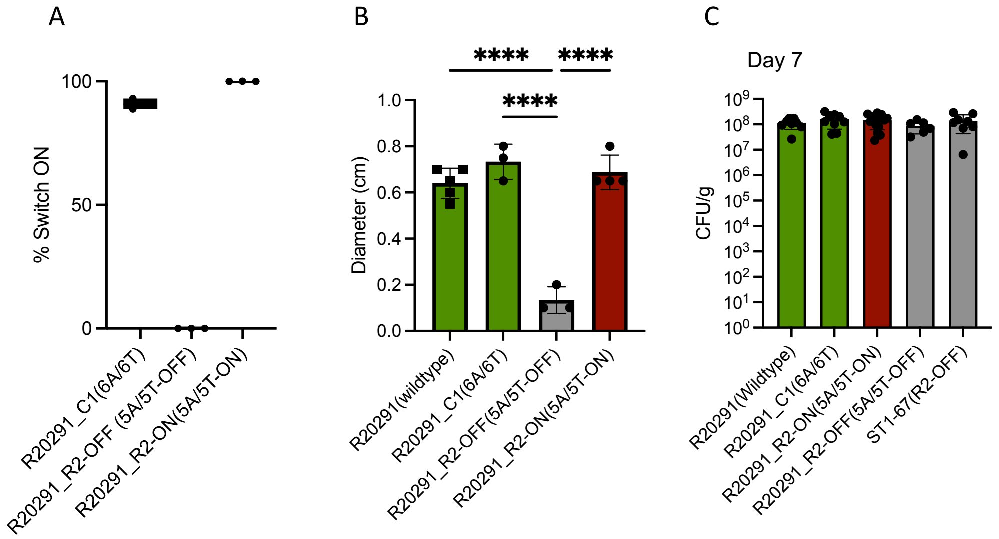


**Figure S4**. The flagellar repeat variants and flagellar directions impact *C. difficile* virulence.
(A) % Flagellar ON vs OFF cells in liquid broth during log phase by qPCR. (B) Swim plates results of *C. difficile* mutants after 24 hours incubation (n = 3 replicates per isolate). (C) Fecal colony-forming units were measured by plating on selective agar on 7 days post-infection. Statistical significance was calculated by One-way ANOVA, **** $p < 0.0001$.

**Table S1**

| Isolate | A-T track in LI | A-T track in LI | IR type | Dominant dire | Virulence |
|---|---|---|---|---|---|
| ST1-11 | 6A-6T | 6A-6T | C1 | ON | high |
| ST1-49 | 6A-6T | 6A-6T | C1 | ON | high |
| ST1-12 | 6A-6T | 6A-6T | C1 | ON | high |
| ST1-53 | 6A-6T | 6A-6T | C1 | ON | high |
| ST1-62 | 6A-6T | 6A-6T | C1 | ON | high |
| ST1-58 | 6A-6T | 6A-6T | C1 | ON | high |
| ST1-26 | 6A-5T | 5A-6T | C2 | ON | high |
| ST1-63 | 6A-5T | 5A-6T | C2 | OFF | intermediate |
| ST1-5 | 6A-5T | 5A-6T | C2 | OFF | intermediate |
| ST1-69 | 6A-6T | 6A-6T | C1 | ON | intermediate |
| ST1-19 | 6A-5T | 5A-6T | C2 | ON | intermediate |
| ST1-51 | 5A-6T | 6A-5T | R1 | ON | low |
| ST1-57 | 6A-5T | 5A-6T | C2 | OFF | low |
| ST1-25 | 6A-5T | 5A-6T | C2 | OFF | low |
| ST1-10 | 6A-5T | 5A-6T | C2 | OFF | low |
| ST1-65 | 6A-5T | 5A-6T | C2 | ON | low |
| ST1-23 | 6A-5T | 5A-6T | C2 | OFF | low |
| ST1-27 | 6A-5T | 5A-6T | C2 | OFF | low |
| ST1-67 | 5A-5T | 5A-5T | R2 | OFF | low |
| ST1-68 | 5A-6T | 6A-5T | R1 | OFF | low |
| ST1-6 | 6A-5T | 5A-6T | C2 | OFF | low |
| ST1-20 | 6A-5T | 5A-6T | C2 | OFF | low |
| ST1-35 | 6A-6T | 6A-6T | C1 | ON | avirulent (cdtR mutant) |
| ST1-75 | 6A-6T | 6A-6T | C1 | ON | avirulent (cdtR mutant) |
| ST1-1 | 6A-6T | 6A-6T | C1 | ON | n/a |
| ST1-2 | 6A-6T | 6A-6T | C1 | ON | n/a |
| ST1-3 | 6A-6T | 6A-6T | C1 | ON | n/a |
| ST1-4 | 6A-6T | 6A-6T | C1 | ON | n/a |
| ST1-7 | 6A-6T | 6A-6T | C1 | ON | n/a |
| ST1-13 | 5A-6T | 6A-5T | R1 | OFF | n/a |
| ST1-14 | 6A-6T | 6A-6T | C1 | ON | n/a |
| ST1-17 | 6A-6T | 6A-6T | C1 | ON | n/a |
| ST1-18 | 6A-6T | 6A-6T | C1 | ON | n/a |
| ST1-21 | 6A-6T | 6A-6T | C1 | ON | n/a |
| ST1-22 | 6A-6T | 6A-6T | C1 | ON | n/a |
| ST1-24 | 6A-6T | 6A-6T | C1 | ON | n/a |

| | | | | | |
|---|---|---|---|---|---|
| ST1-28 | 6A-6T | 6A-6T | C1 | ON | n/a |
| ST1-29 | 6A-6T | 6A-6T | C1 | ON | n/a |
| ST1-32 | 6A-5T | 5A-6T | C2 | OFF | n/a |
| ST1-33 | 6A-5T | 5A-6T | C2 | OFF | n/a |
| ST1-34 | 6A-6T | 6A-6T | C1 | ON | n/a |
| ST1-36 | 6A-6T | 6A-6T | C1 | ON | n/a |
| ST1-37 | 6A-5T | 5A-6T | C2 | OFF | n/a |
| ST1-38 | 6A-6T | 6A-6T | C1 | ON | n/a |
| ST1-39 | 6A-6T | 6A-6T | C1 | ON | n/a |
| ST1-40 | 6A-6T | 6A-6T | C1 | ON | n/a |
| ST1-41 | 6A-5T | 5A-6T | C2 | OFF | n/a |
| ST1-42 | 6A-6T | 6A-6T | C1 | ON | n/a |
| ST1-45 | 6A-5T | 5A-6T | C2 | OFF | n/a |
| ST1-46 | 6A-6T | 6A-6T | C1 | ON | n/a |
| ST1-47 | 6A-6T | 6A-6T | C1 | ON | n/a |
| ST1-48 | 6A-6T | 6A-6T | C1 | ON | n/a |
| ST1-50 | 6A-6T | 6A-6T | C1 | ON | n/a |
| ST1-52 | 6A-5T | 5A-6T | C2 | OFF | n/a |
| ST1-54 | 6A-6T | 6A-6T | C1 | ON | n/a |
| ST1-55 | 6A-6T | 6A-6T | C1 | ON | n/a |
| ST1-59 | 6A-5T | 5A-6T | C2 | OFF | n/a |
| ST1-66 | 6A-6T | 6A-6T | C1 | ON | n/a |
| ST1-70 | 6A-6T | 6A-6T | C1 | ON | n/a |
| ST1-71 | 5A-6T | 6A-5T | R1 | OFF | n/a |
| ST1-72 | 6A-6T | 6A-6T | C1 | ON | n/a |
| ST1-73 | 6A-5T | 5A-6T | C2 | OFF | n/a |
| ST1-79 | 6A-6T | 6A-6T | C1 | ON | n/a |
| ST1-31 | 6A-5T | 5A-6T | C2 | OFF | n/a |

### Table S2. Oligos used in this study for CRISPR mutagenesis.

| Name | Sequence 5'-3' | Reference | Purpose |
|---|---|---|---|
| R_FlgSwitch_up_For | AAACAGCTATGACCGCGGCCGCC ATTTATTTAAATTTCGTTAATTGAC | This study | Forward primer amplifying upstream region of flagellar switch; also for amplifying out flagellar switch region from *C. difficile* isolates with IR variants. |
| R_FlgSwitch_up_Rev | AACTTTCAGTTTAGCGGTCTGGGCGCCGG TATAATATGTTATTATAGGTGTTTTTTAC | This study | Reverse primer amplifying upstream region of flagellar switch. |
| R_FlgSwitch_down_For | GGCGCCCAGACCGCTAAACTGAAAGT TAAATATATTTTGAGGAGGAGTGAGT | This study | Forward primer amplifying downstream region of flagellar switch. |
| R_FlgSwitch_down_Rev | TTATTTTTATGCTAGCTCGAGGTTTTC TTCAAATGAAACAACTTTTCT | This study | Reverse primer amplifying downstream region of flagellar switch; also for amplifying out flagellar switch region from *C. difficile* isolates with IR variants. |
| R_FlgSwitch_sgRNA1_For | AATTAAACTGTAAATGGCCAAATATAATAATG TATCACTTGTTTTAGAGCTAGAAATAGC | This study | Forward primer gRNA Cloning into pCE677 digested with MscI and MluI, targeting flagellar switch. |
| R_RFP_sgRNA2_For | AATTAAACTGTAAATGGCCACAGTTTAGCGGTCTG GGCGCGTTTTAGAGCTAGAAATAGC | This study | Forward primer gRNA Cloning into pCE677 digested with MscI and MluI, targeting RFP landing pad. |
| R_RFP_sgRNA3_For | AATTAAACTGTAAATGGCCACCTATAATAACAT ATTATACGTTTTAGAGCTAGAAATAGC | This study | Forward primer gRNA Cloning into pCE677 digested with MscI and MluI, targeting RFP landing pad. |
| CDEP4427 | ATAGTTGCAGAGCTTACGCGT CTAGTCAGACATCATGCTGAT | [38] | Universal Reverse primer for gRNA Cloning into pCE677 digested with MscI and MluI |

**Table S3. Plasmids used in the *E. coli* colorimetric assay.**

| Name | Description | Replication origin | Resistance | Reference |
|---|---|---|---|---|
| pQD1 | pBAD derivative for RecV expression | p15a | chloramphenicol | This study |
| pQD2 | C1 invertible promoter 6A/6T – OFF (red) | pSC101 | kanamycin | This study |
| pQD3 | C1 invertible promoter 6A/6T – ON (green) | pSC101 | kanamycin | This study |
| pQD4 | C2 invertible promoter 5A/6T – OFF (red) | pSC101 | kanamycin | This study |
| pQD5 | C2 invertible promoter 5A/6T – ON (green) | pSC101 | kanamycin | This study |
| pQD6 | R2 invertible promoter 5A/5T – OFF (red) | pSC101 | kanamycin | This study |
| pQD7 | R2 invertible promoter 5A/5T – ON (green) | pSC101 | kanamycin | This study |
